# Deciphering shared and divergent tissue architectures from cross-species spatial transcriptomics

**DOI:** 10.64898/2026.06.16.732760

**Authors:** Biao Zhang, Xiang Zhou, Shuqin Zhang, Shihua Zhang

## Abstract

The integration of spatial transcriptomics (ST) data across species is essential for cross-species and translational studies, but remains challenging due to molecular divergence and anatomical differences between organisms. We present STACAME, a graph attention autoencoder-based framework to decipher shared and divergent tissue architectures from cross-species ST data by explicitly modeling both orthologous and species-specific genes. STACAME aligns ST slices in a spatially aware manner, identifies homologous and species-specific domains, and enables a suite of downstream comparative analyses. We demonstrate its utility by integrating ST datasets from diverse tissues, including hippocampus, isocortex, embryo, breast, liver, and cerebellum, across multiple species such as human, macaque, marmoset, mouse, and zebrafish. STACAME supports cross-species spatial domain alignment, the detection of shared and divergent spatially variable genes, development alignment and comparison, and the 3D integration of tissue architecture. This flexible approach facilitates the translation of findings from model organisms to humans, providing a unified computational platform for cross-species spatial transcriptomics.

## Introduction

Cross-species comparative analysis has been a cornerstone of evolutionary biology and human disease research since the era of Darwin^1–3^. It enables systematic investigation of functional conservation and divergence across species, illuminating the evolutionary origins of morphological and functional traits^4–8^. By comparing gene regulation and cellular behaviors across organisms^9–11^, such analyses reveal the evolutionary pressures shaping diverse biological systems, while also allowing discoveries from model organisms, such as mice and macaques, favored for their short life cycles, ease of husbandry, and genetic tractability, to be translated to human biology^12–14^.

Recent advances in ST technologies, which jointly measure messenger RNA expression and spatial organization within tissues, now provide an unprecedented opportunity for cross-species comparisons. However, despite the growing dominance of non-human spatial transcriptomics (ST) datasets^7,15,16^, transferring insights to humans at the whole-tissue level remains challenging due to substantial evolutionary divergence, including large-scale anatomical differences such as the expansion of the human cortex^17,18^, as well as historically inconsistent terminology used to describe homologous tissues across species. This underscores the critical need to integrate ST data from diverse species into a unified spatial framework and to align corresponding tissue domains, thereby facilitating robust transfer of biological insights across species, including to human systems.

Various ST technologies have been developed with different spatial resolutions^17–20^, which have facilitated critical research, including characterizing the biological architecture of tissues^20,22^, identifying biomarkers of complex brain diseases^23^, and understanding the development process of organogenesis^21^. In particular, computational methods have been successfully applied not only to single-condition ST data^24–26^, but also to the integration of ST data across diverse conditions, technologies, or developmental stages^27–30^, supporting the identification of spatially variable genes^31,32^, characterization of the tissue structures, and deciphering spatial cell-cell communication patterns^33^ more conveniently. As high-resolution ST data of various species, such as the whole brain cortex or embryos of mice, macaques, and humans^34–38^, emerge, an essential problem is how to effectively integrate these ST data into a unified representation space to enable cross-species comparative analysis. However, current ST data integration methods are not designed to address interspecific differences, such as unaligned gene expression vectors that blend with data heterogeneity introduced by variations of tissue organization, profiling technologies, experimental conditions, and developmental speed^10,39^.

Existing methods for molecular-level alignment studies, such as single-cell integration approaches, have mostly focused on addressing cross-species homologies through investigations of cell type commonality and diversity across species^9,39–41^, without modeling spatial coordinates. Thus, these methods, such as Harmony^42^, are not proper for cross-species ST data integration. For the recently developed spatial integration approaches, such as SEDR^43^, PASTE^44^, SPIRAL^28^, Nichformer^30^, and Novae^45^, their performance on cross-species integration tasks has not yet been tested.

For these methods, only one-to-one gene orthologs are used, because the model was originally designed for integrating RNA-seq data or ST slices of the same species. For example, STAligner^27^ utilized a graph attention auto-encoder to integrate spots from different slices, and the integration of mouse and human embryos demonstrated that STAligner facilitates cross-species integration on one-to-one gene orthologs. The many-to-many gene orthologs have been proven beneficial for bridging cross-species genomics, especially for evolutionarily distant species.

To overcome these challenges, we developed STACAME, a novel framework that integrates spatial transcriptomic data across diverse species through a graph neural network-based approach. We demonstrate STACAME’s versatility and robustness by applying it to integrate ST data from multiple species across diverse biological contexts, including hippocampus, isocortex, and cerebellum across mice, marmosets, and macaques; embryonic development across mice, zebrafish, and humans; and disease models, such as Alzheimer’s disease and breast cancer. This framework provides a powerful tool for cross-species and translational research, enabling a more comprehensive understanding of biological processes across species.

## Results

### Overview of STACAME

STACAME employs a three-stage pipeline to integrate spatial transcriptomics data across species within a unified graph framework: (i) ortholog-based graph construction, (ii) species-specific pretraining, and (iii) multi-species alignment (**Fig. 1a**). It first exhaustively traverses many-to-many homologous gene combinations to build equal-length feature vectors, enabling direct cross-species comparison and the construction of a joint graph that connects spatial neighbors within each species to cross-species spot correspondences. In the pretraining stage, a graph attention autoencoder learns species-specific embeddings independently for each species, using only intra-species spatial edges; this isolates species-specific spatial domains before any cross-species mixing and provides a stable initialization. The alignment stage then concatenates these embeddings and feeds them into a second graph attention autoencoder that projects all spots into a shared latent space.

**Figure 1:**
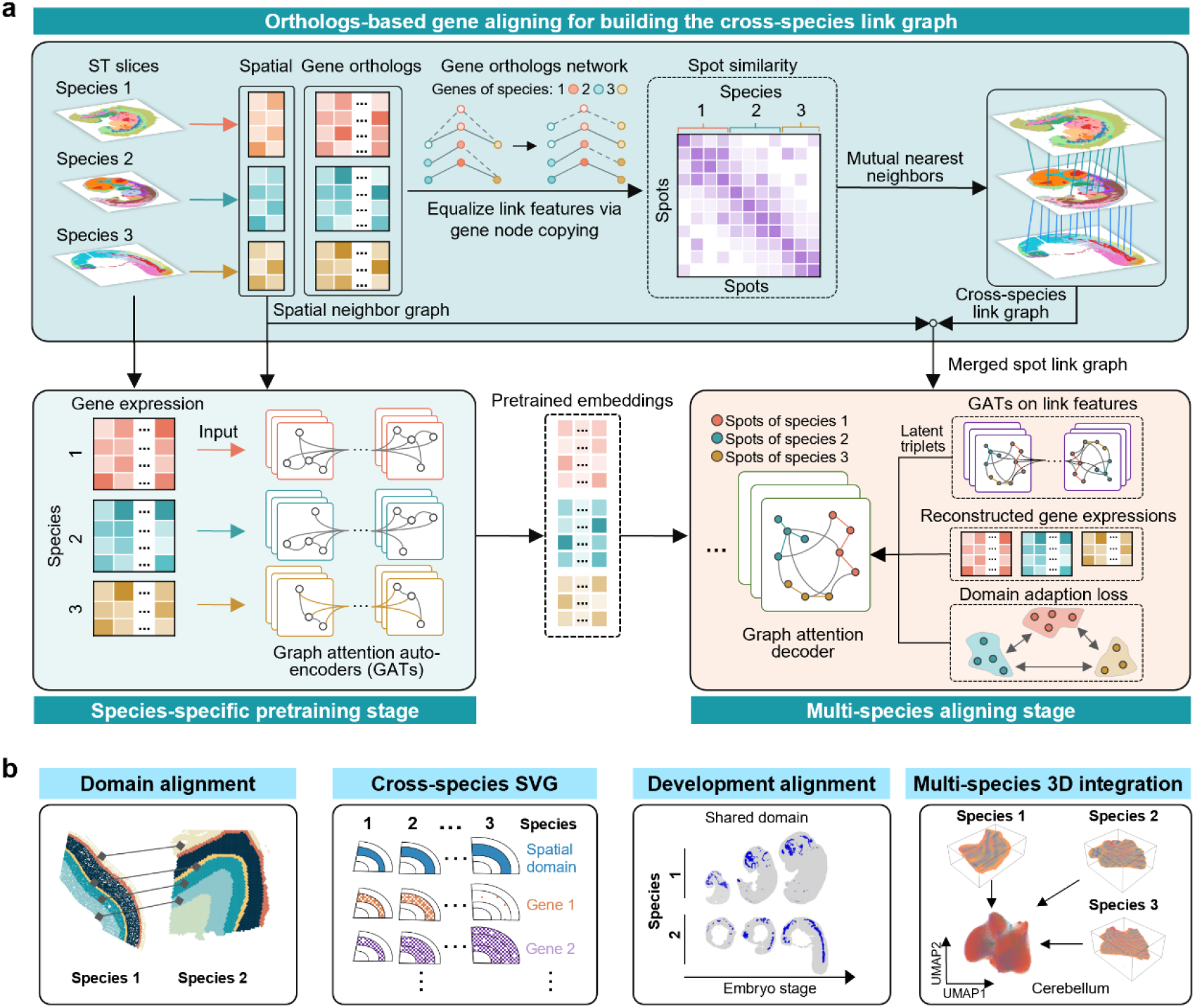
Overview of STACAME and its applications. **a**, Workflow of STACAME. In the orthologs-based gene aligning stage for building the cross-species link graph, spatial neighbor networks are constructed within each species using Euclidean distances between spot coordinates. An expression matrix of equal gene length (*X*_*e*_) is generated by mapping orthologous genes across species. Cross-species spot correspondences are then initially inferred using either the mutual nearest neighbors (MNN) algorithm or, optionally, k-nearest neighbors (k-NN). In the species-specific pretraining stage, homologous and species-specific highly variable genes (HVGs) are selected as input features for a graph attention network (GAT), where the graph topology is defined by the intra-species spatial neighbor network. In the multi-species aligning stage, pretrained embeddings are aligned and decoded through another GAT that incorporates cross-species edges to reconstruct gene expression (optimized via mean squared error loss). Cross-species triplets are constructed by identifying inter-species MNN spot pairs in both the gene expression space and the latent space, where the latent space is generated by an auxiliary lightweight graph attention autoencoder using the PCA of *X*_*e*_. Maximum mean discrepancy (MMD) and adversarial (GAN) losses are applied to facilitate domain adaptation. **b**, Applications enabled by STACAME include cross-species domain alignment, identification of cross-species shared or divergent spatially variable genes, comparative analysis of organ development across species, and three-dimensional (3D) integration of ST data from multiple species.

Alignment is driven by a pair of complementary triplet losses that work at different resolutions. An initial triplet loss relies on mutual nearest neighbors (MNN) identified directly in the original expression space to establish coarse cross-species correspondences. However, when species heterogeneity is high, such expression-space pairs can be noisy. To overcome this, an auxiliary triplet loss leverages a lightweight graph attention autoencoder trained on the ortholog-aligned gene vectors; by dynamically recomputing MNN pairs in its learned latent space, it iteratively refines anchor-positive relationships, capturing subtle homology-driven similarities that would otherwise be missed. This dual-loss strategy is key to robustly aligning divergent species. Maximum mean discrepancy (MMD) and adversarial domain adaptation losses then jointly harmonize global distributions. MMD aligns statistical moments while an adversarial discriminator removes complex, higher-order discrepancies. In addition, the manifold-preserving loss, weighted by pretrained embedding affinities, safeguards the intrinsic neighborhood topology of each species, ensuring that biologically meaningful local structures are not distorted during alignment.

We applied STACAME to processed gene expression data and spatial coordinates of spots from humans, macaques, mice, and zebrafish, covering diverse healthy and pathological organs sequenced on various platforms, including 10x Visium, Stereo-seq, Slide-seqV2, Slide-tags, 10x Xenium, and MERFISH. By mapping cross-species spatial spots into a shared embedding space, STACAME enables the identification of cross-species conserved spatial domains, the discovery of shared or divergent spatially variable genes, and comparative analyses of tissue development and disease mechanisms (**Fig. 1b**).

### STACAME improves cross-species spatial domain alignment of ST data

We applied STACAME to the macaque^37^ and human^22^ dorsolateral prefrontal cortex (DLPFC) data produced by Stereo-seq and 10x Visium, respectively (**Fig. 2a**). With STACAME, the low-dimensional embeddings of spots for both species are obtained. The UMAP (uniform manifold approximation and projection) of the latent representations from different methods is presented in **Fig. 2b**. STACAME achieved higher levels of biological conservation, species mixing, and other separate metrics compared to Harmony, Scanorama, SEDR, and STAligner, which only use one-to-one homologous genes (**Fig. 2c**; **Supplementary Fig. 1a**). Although almost all methods mix spots from both species, only STAligner and STACAME align corresponding cortex layers and identify human white matter as an independent domain. To test whether STACAME could identify cortex layer domains in both species’ datasets, we ran *mclust*^46^ (clustering number = 7) on the integrated embeddings and computed the overall ARI (adjusted rand index) between manually annotated layers and clustered domains. The results show that STACAME accurately identifies spatial domains aligned with manually annotated cortex layers, achieving a higher ARI than other methods (**Fig. 2d**). Compared to STAligner, STACAME distinguishes human white matter from other cortex layers. The Sankey plot of alignment among macaque DLPFC layers, common clusters, and human DLPFC layers (with line width indicating the proportion of spot numbers) also depicts that STACAME identifies common domains shared between macaque and human (**Fig. 2e, Supplementary Fig. 1b**). Compared to other methods, as shown in **Supplementary Fig. 1c**, the homologous regions’ embeddings produced by STACAME show higher significance (*P*-value < 10^−3^).

**Figure 2:**
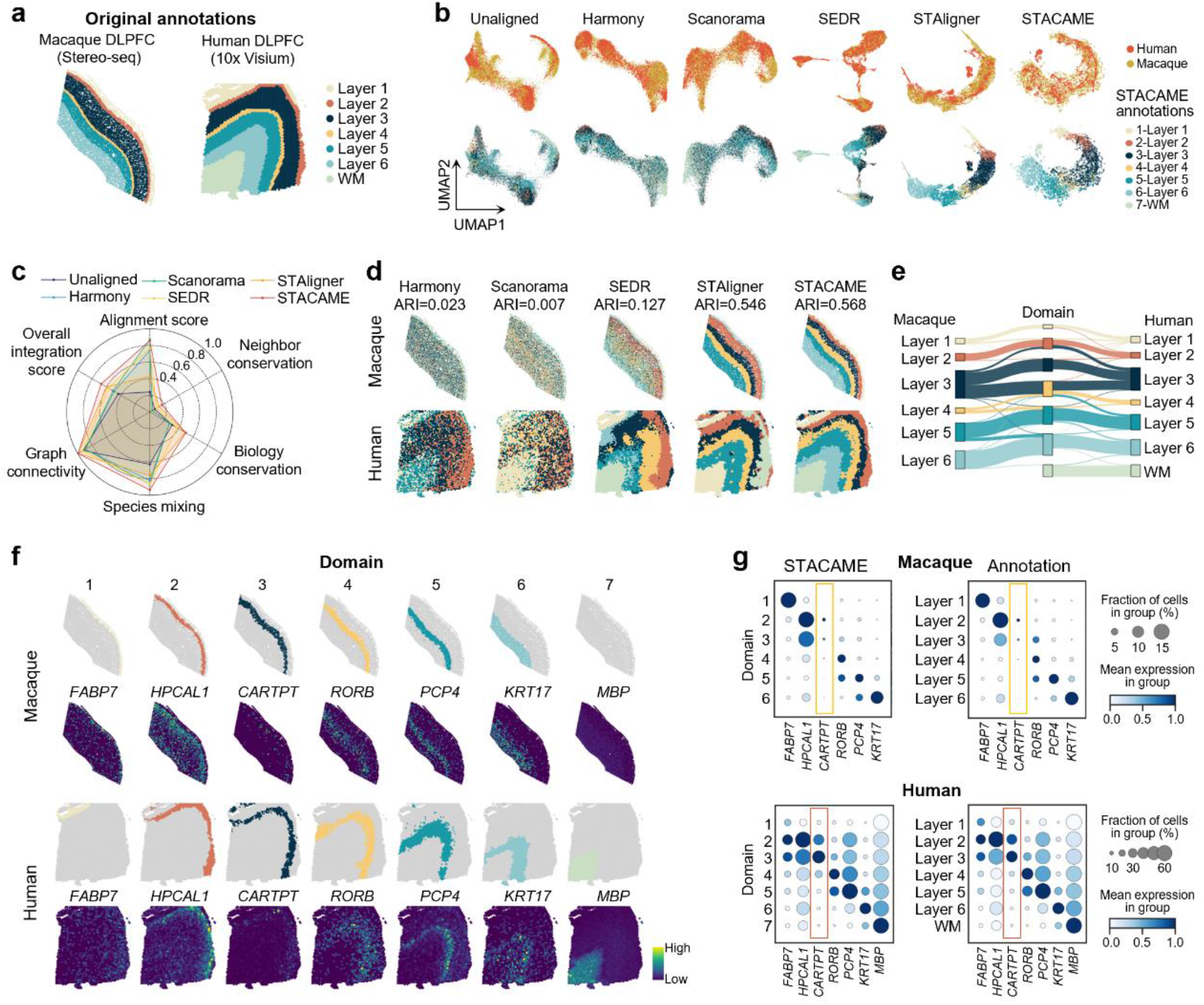
Cross-species alignment of dorsolateral prefrontal cortex (DLPFC) spots between macaques and humans. **a**, Manually annotated laminar labels in macaque (Stereo-seq)^37^ and human (10x Visium) DLPFC^22^. **b**, UMAP visualization of spot embeddings generated by multiple integration methods. **c**, Quantitative comparison of integration performance using multiple evaluation metrics. **d**, Spatial domains identified by STACAME and baseline methods. *mclust* (k = 7) was applied to integrated embeddings, and the adjusted Rand index (ARI) was computed between clustering results and manual layer annotations. **e**, Sankey diagram showing alignments among macaque DLPFC layers, shared clusters, and human DLPFC layers; line width reflects the proportion of spots. **f**, Spatial maps of identified DLPFC layer domains and their marker genes. Human markers are from prior literature; macaque markers are one-to-one orthologs of human genes. **g**, Dot plots showing expression levels (color) and detection frequencies (size) of marker genes in macaque and human domains (left) and annotated layers (right). The right panels show spot counts per group. Differences in expression are assessed by Wilcoxon rank-sum test.

To validate the rationale of cross-species shared domains, we visualized the spatial expression of human cortex layer marker genes (layers 1 to 6 and white matter) and the homologous counterparts in the macaque cortex, alongside the corresponding domains identified by STACAME. As shown in **Fig. 2f**, most marker genes and the common domains display a high level of overlap. The dot plots of differential expression reveal that *CARTPT* has significantly higher specificity in human layer 2 than in macaque layer 2 (**Fig. 2g**). The expression levels and detection frequencies of marker genes are quite similar between domains (left) and annotated layers (right) in both species, including the divergent layer 3 marker gene *CARTPT* (**Fig. 2g**), indicating that the shared domains identified by STACAME are accurate.

We next benchmarked STACAME across a diverse array of cross-species integration challenges, encompassing distinct brain regions, sequencing platforms, and phylogenetic distances. When integrating Stereo-seq data from mouse, marmoset and macaque hippocampus, STACAME consistently yielded superior scores in biological conservation, species mixing and overall integration (**Supplementary Fig. 2a, b**). Visualization of the embeddings confirmed thorough interspecies mixing with preserved separation of homologous anatomical substructures (**Supplementary Fig. 2c**). This accurate alignment was supported by Sankey diagrams (**Supplementary Fig. 2d**) and quantitatively confirmed by significantly higher correlations (*P* < 10^−3^) between embeddings of spots from homologous regions versus random pairs (Welch’s t-test with Bonferroni correction; **Supplementary Fig. 2e**).

STACAME demonstrated robustness to technical variation in a cross-platform integration of mouse (Slide-seqV2) and macaque (Stereo-seq) hippocampus data. The derived clusters showed the highest concordance with manually annotated hippocampal subregions (Adjusted Rand Index; **Supplementary Fig. 3c**) and outperformed other methods on all component metrics (**Supplementary Figs. 3b** and **4a**). Marker gene expression patterns validated the biological identity of the integrated domains (**Supplementary Fig. 3d**). Embeddings were well-mixed across species (**Supplementary Fig. 4b**), with clear alignment of homologous areas (**Supplementary Fig. 4c**) and significantly enhanced within-area correlations (**Supplementary Fig. 4d**).

STACAME also effectively scaled to integrate multiple biological replicates, simultaneously aligning three spatial slices each from the mouse and macaque hippocampus. It achieved the highest overall integration score (**Supplementary Fig. 5a, b**) and the best agreement between derived clusters and expert annotations (**Supplementary Fig. 5c**), while maintaining well-mixed embeddings that preserved intra-species spatial structure (**Supplementary Fig. 5d**).

Finally, applied to MERFISH profiles of mouse and human cortical layers, STACAME achieved the highest overall integration score (**Supplementary Fig. 6b**), well-integrated embeddings (**Supplementary Fig. 6c**), and clusters that most closely matched the annotated laminar architecture (**Supplementary Fig. 6d**). Together, these results demonstrate that STACAME enables robust and accurate integration of cross-species ST data across a wide range of tissues, technologies, and experimental designs.

### STACAME reveals cross-species shared and divergent spatially variable genes

One fundamental application of cross-species ST comparison is to identify shared spatial domains and characterize the underlying conserved and divergent transcriptional genes. We first applied STACAME to integrate hippocampal datasets from mice, marmosets, and macaques generated using Stereo-seq. STACAME achieved superior integration performance compared to baseline methods (**Supplementary Fig. 2**) and accurately mapped hippocampal spots from different species into a shared embedding space. Clustering of the integrated embeddings using *mclust* (k = 9) revealed well-defined hippocampal subregions, including CA1, CA2, CA3, and dentate gyrus, which were highly consistent with manual annotations (**Fig. 3a**).

**Figure 3:**
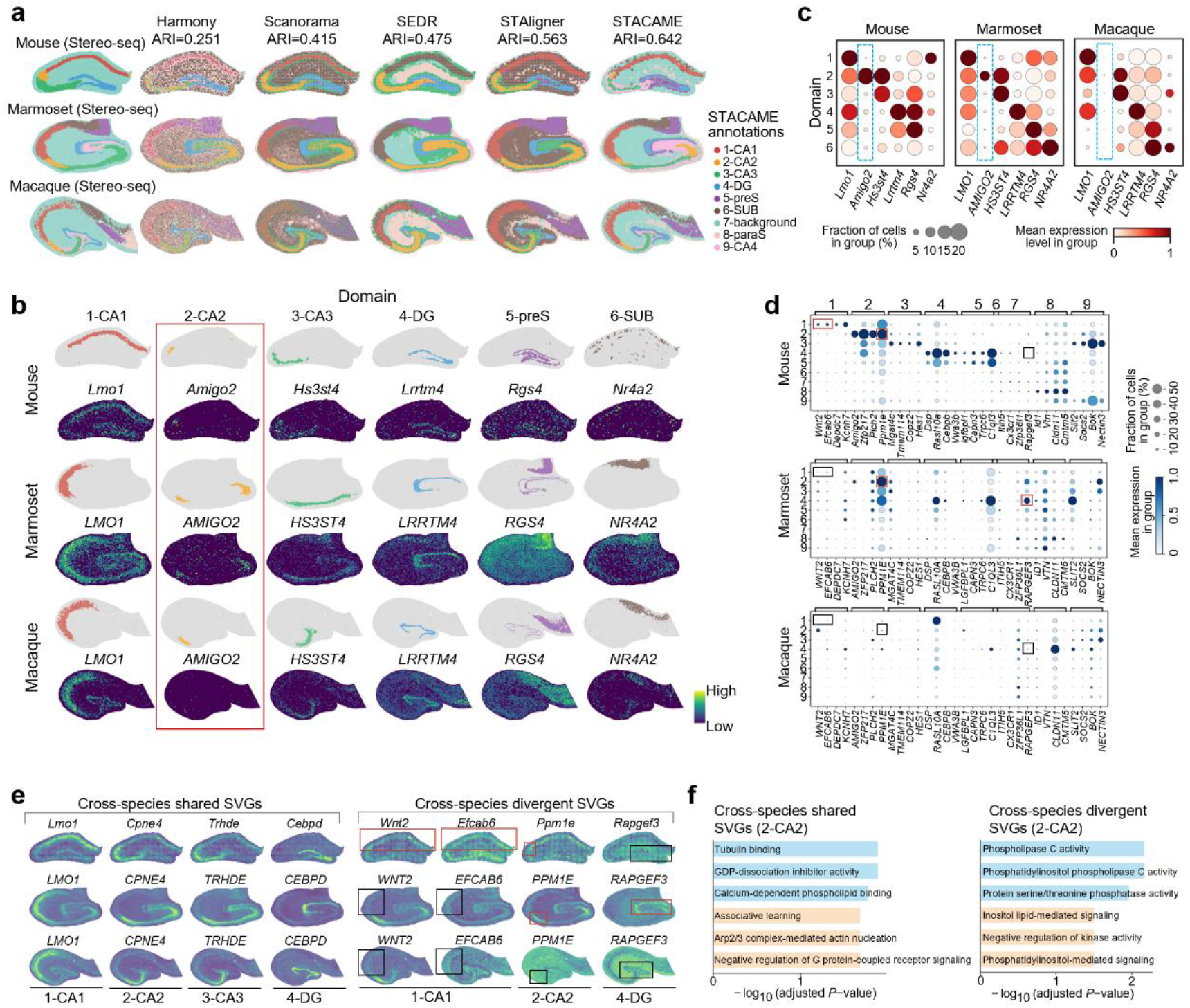
Cross-species shared and divergent spatially variable genes in the hippocampus of mouse, marmoset, and macaque. **a**, Spatial domains identified by STACAME and baseline methods. *mclust* (k = 9) was applied to integrated embeddings, and ARI was calculated against manual annotations. **b**, Spatial maps of hippocampal domains and associated marker genes. Mouse markers are derived from the literature, and marmoset and macaque markers are one-to-one orthologs of the corresponding mouse genes. **c**, Dot plots of marker gene expression (color) and detection frequency (size) across CA1, CA2, CA3, and dentate gyrus domains in all three species. The right panels indicate spot numbers. Notably, the macaque ortholog of the mouse CA2 marker *Amigo2* shows no significant differential expression in macaque CA2. **d**, Cross-species divergent spatially variable genes (CDSVGs): mouse-specific differentially expressed genes (DEGs) whose orthologs lack significant differential expression in marmoset or macaque, e.g., *Wnt2*, Efcab6, Ppm1e, and Rapgef3, *Amigo2* (highlighted by frame, red frame: significant, black frame: not significant). **e**, Spatial expression patterns of representative cross-species shared (left) and divergent (right) spatially variable genes using STACAME reconstructed gene expressions. The frame indicate the corresponding domain of mouse marker genes (red frame: significant, black frame: not significant). **f**, Gene set enrichment analysis (GSEA) for cross-species shared and divergent genes relative to mouse hippocampal domain 2. Enrichment analysis was carried out with Enrichr (via GSEApy) against the ‘Mouse’ organism using the gene sets GO_Biological_Process_2023, GO_Cellular_Component_2023, GO_Molecular_Function_2023 and Mouse_Gene_Atlas. Bar length represents − log_10_(adjusted *P* value) from Fisher’s exact test; the top three terms by adjusted *P* value are shown for each gene set.

To investigate the conserved and divergent spatial gene expression patterns, we visualized the spatial expression of canonical mouse hippocampal marker genes and their one-to-one orthologs in marmoset and macaque across CA1, CA2, CA3, and dentate gyrus domains identified by STACAME (**Fig. 3b**). Notably, the well-established mouse CA2 marker *Amigo2*^47^ does not exhibit significantly larger differential expression field in marmoset or macaque CA2, and. In addition, the differential expression patterns of *Lmo1, Hs3st4*, and *Lrrtm4* are consistently focused on known hippocampal regions among the three species. The pre-subiculum marker gene *Rgs4* and subiculum marker gene *Nr4a2* exhibit a more spatially confined expression pattern on corresponding STACAME domains for marmoset and macaque than mouse, consistent with the fact that pre-subiculum and subiculum are not identified by STAGATE for the mouse slice. Dot plot analyses of row-normalized gene expression confirmed the above findings (**Fig. 3c**). Although *AMIGO2* is robustly enriched in mouse CA2, it is expressed in a markedly smaller fraction of spots, lacking clear spatial specificity in macaque CA2. Besides, unlike marmoset and macaque, *Nr4a2* shows differential expression in the mouse domain 1-CA1 instead of 6-SUB, probably due to the lack of subiculum in the mouse slice. Independent validation using Allen Mouse Brain Atlas ISH data and Human Protein Atlas protein expression data excluded technical artifacts or data quality issues as potential explanations (**Supplementary Fig. 7a, b**). These results suggest that the spatial expression domain of *Amigo2* is not evolutionarily conserved and does not scale with the expansion of CA2 size from mouse to primates.

To characterize divergent genes across species, we compared CA2 domains from mice, marmosets and macaques. Amigo2 was significantly upregulated in the mouse CA2 but not in the macaque CA2 (**Supplementary Fig. 7c**). We further identified top significant (measured by adjusted *P*-values with positive logarithm of fold change) mouse-specific differentially expressed genes (DEGs) whose orthologs lacked significant differential expression (adjusted *P*-values > 0.05) in marmoset or macaque, defining a set of cross-species divergent spatially variable genes (CDSVGs; **Fig. 3d**). Representative CDSVGs, including *Wnt2, Efcab6, and Ppm1e*, showed spatially restricted expression in mice but not in macaques or marmosets (**Fig. 3e**; **Supplementary Fig. 7c-e**). The mouse marker gene of domain 7, Rapgef3, however, was significantly enriched in domain 4-DG of the marmoset, presenting a typical example of CDSVGs and highlighting the existence and complexity of species-specific spatial expression divergence. Conversely, cross-species conserved spatially variable genes (CCSVGs), which are filtered out by selecting homologous genes with top adjusted *P*-values and positive logarithm of fold change for three species, exhibited consistent domain-specific expression across all three species (**Fig. 3e**; **Supplementary Fig. 7f**). Gene set enrichment analysis revealed that the conserved transcriptional programs were primarily associated with fundamental neuronal and hippocampal functions, including cytoskeletal organization and synaptic processes (for example, tubulin binding^48^, GDP-dissociation inhibitor activity, calcium-dependent phospholipid binding, and associative learning). In contrast, cross-species divergent programs were enriched for signaling and regulatory pathways implicated in species-specific neuronal differentiation and functional specialization, including phospholipase C activity^49^, phosphatidylinositol- and inositol lipid–mediated signaling, and negative regulation of kinase and G protein–coupled receptor signaling pathways (**Fig. 3f**).

In short, STACAME can reveal that homologous marker gene expression patterns varied substantially across species despite anatomical correspondence (**Supplementary Fig. 8**), underscoring the importance of cross-species integration of ST data. These results demonstrate that STACAME enables systematic identification of homologous spatial domains while disentangling shared and species-specific transcriptional programs, providing a powerful framework for comparative studies of brain evolution and function.

### STACAME enables robust identification of shared and divergent tissue structures across phylogenetically distant species

Integrating ST data across phylogenetically distant species remains challenging because the proportion of one-to-one homologous genes decreases substantially with evolutionary distance^10,39,50^. To evaluate the ability of STACAME to resolve shared and divergent tissue architectures under such conditions, we first performed a stage-resolved integration of mouse and zebrafish embryonic tissues across multiple developmental time points (**Fig. 4**). Specifically, we integrated mouse embryos at stages E9.5, E11.5 and E12.5 and zebrafish embryos at stages 12hpf, 18hpf and 24hpf, all generated using Stereo-seq. The embeddings of all spots from the six developmental stages obtained using STACAME were jointly clustered using *mclust* (k = 15), yielding spatial domains shared across species and stages (**Fig. 4a**; **Supplementary Fig. 9a, b**). Many shared domains corresponded to conserved anatomical structures, including forebrain, hindbrain, muscle and cartilage primordium, with spatial organization and marker gene expression patterns that were highly consistent across species (**Supplementary Fig. 9c**). Among the shared domains, five domains were consistently associated with brain-related tissues across both species, as confirmed by their spatial distributions and the expression of canonical marker genes (**Fig. 4b, c**). The proportions of spots assigned to each brain-relevant domain varied systematically across developmental stages, reflecting conserved yet dynamic tissue composition during embryogenesis (**Fig. 4d**). Notably, STACAME also identified biologically meaningful species-specific domains that only exist in one species, such as a mouse-specific domain marked by *Sox9* (and its zebrafish homolog *sox9a*), whose spatial localization and functional enrichment were consistent with known developmental roles (**Fig. 4e, f**).

**Figure 4:**
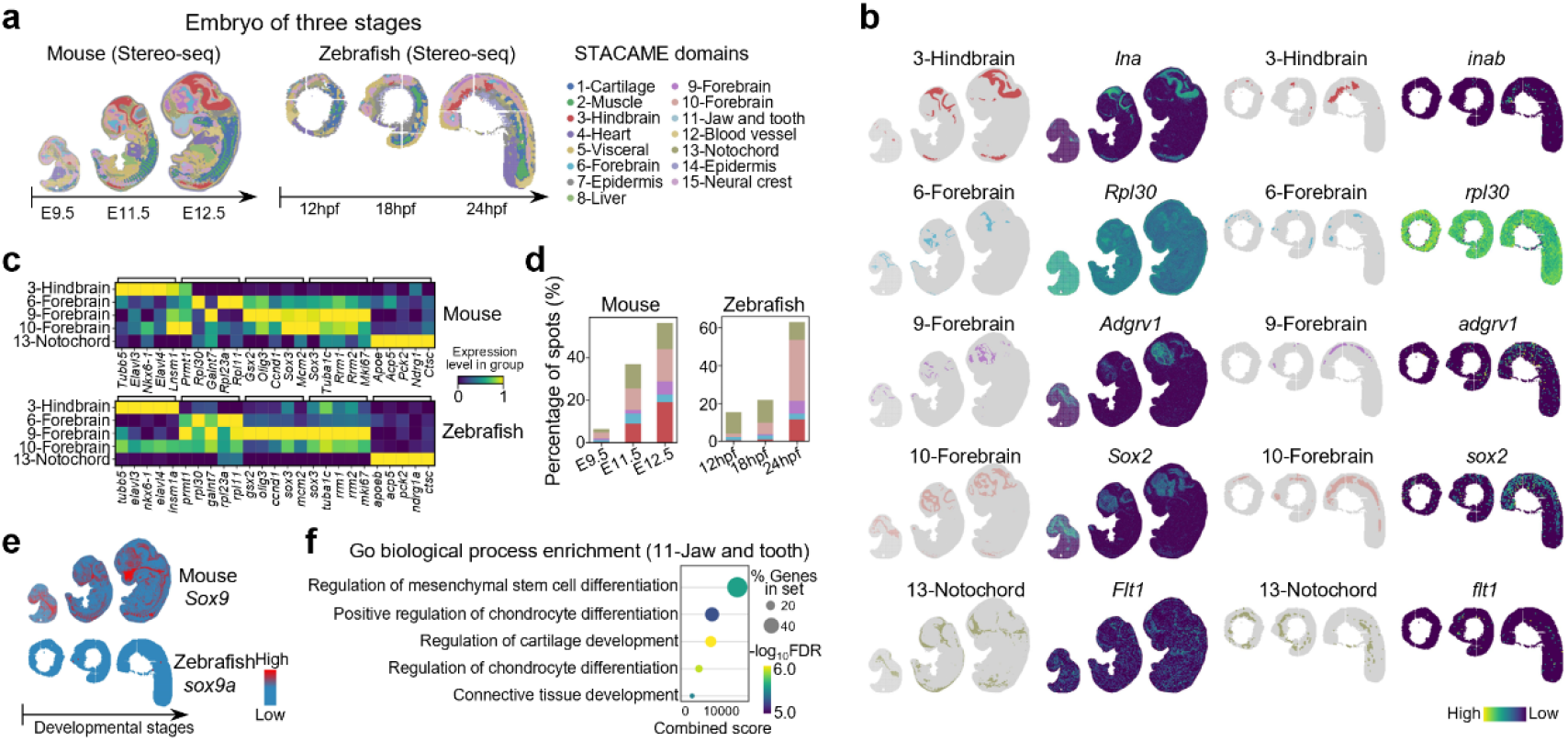
Integration of phylogenetically distant embryonic tissues across developmental stages. Mouse embryo at stage E9.5, E11.5 and E12.5 (Stereo-seq)^21^ and zebrafish embryo at stage 12hpf, 18hpf and 24hpf (Stereo-seq)^78^. **a**, 15 domains identified by the *mclust* method on the STACAME spot embeddings of multiple mouse (E9.5, E11.5 and E12.5) and zebrafish (12hpf, 18hpf and 24hpf) embryo slices. **b**, The spatial map of the five brain-relevant species shared domains and their marker genes. **c**, The identified marker genes of five brain-relevant species shared domains. **d**, Proportion of spots assigned to each brain-relevant domain per embryo slice. **e**, Spatial map of one typical marker gene (Sox9 for mouse, sox9a for zebrafish) for domain 11, which indicates that the mouse-specific domain is biologically meaningful. **f**, GSEA results for domain 11. The top 20 domain‐specific marker genes ranked by statistical significance were used for Gene Ontology (GO) biological process enrichment analysis with Enrichr via GSEApy. The dot plot displays the five most significantly enriched terms (adjusted P < 0.05, Benjamini–Hochberg correction). Dot colour encodes the adjusted P value, and dot size represents the number of input genes overlapping each term.

We next extended the analysis to a three-species integration involving human, mouse and zebrafish embryos to assess the scalability of STACAME across greater evolutionary distances (**Supplementary Figs. 10 and 11**). Human embryos at Carnegie stage 13 (10x Visium), mouse embryos at stage E11.5 (Stereo-seq), and zebrafish embryos at stage 24hpf (Stereo-seq) were jointly embedded using STACAME. Mouse anatomical annotations and corresponding marker genes were used as reference standards to evaluate shared domains. Clustering of the integrated embeddings with *mclust* (k = 19, matching the number of annotated mouse tissues) yielded homologous spatial domains across the three species (**Supplementary Fig. 10b, c**). Many shared domains corresponded to conserved anatomical structures, including forebrain, hindbrain, muscle and cartilage primordium, whose spatial organization and marker gene expression patterns remained highly consistent across species. Quantitative analysis further demonstrated strong correspondence between annotated tissues and inferred domains, as reflected by high inter-species Pearson correlations of averaged expression profiles (**Supplementary Fig. 11**).

While most shared tissue structures were robustly identified, certain zebrafish organs, such as the heart, exhibited reduced spatial concordance with their corresponding domains. Further analysis revealed that this discrepancy was attributable to the relatively small spatial extent of the zebrafish heart at 24hpf compared to mouse and human embryos, resulting in limited representation in spatial transcriptomic slices (**Supplementary Fig. 10d-g**). Despite this limitation, heart marker genes such as *hand2* remained highly expressed within the identified corresponding domains, indicating that STACAME captures meaningful molecular signatures even when anatomical structures are underrepresented. Together, these results demonstrate that STACAME enables robust identification of conserved and divergent tissue domains across large evolutionary distances and developmental stages, while also highlighting biologically interpretable sources of cross-species variability.

### STACAME enables robust identification of disease-associated spatial domains across species and disease contexts

We next assessed the capability of STACAME to detect disease-associated spatial domains through cross-species integration of diseased and normal tissues. We integrated a normal macaque hippocampus section profiled by Stereo-seq with a mouse hippocampus section under Alzheimer’s disease (AD) conditions profiled by Slide-seqV2^23^. Clustering of STACAME embeddings using *mclust* identified 15 spatial domains across the two species (**Fig. 5a**). Most domains corresponded to canonical hippocampal subregions and were well aligned between species, including CA1, CA3 and dentate gyrus (**Fig. 5b**). However, two species-specific domains were clearly observed: a mouse-specific domain (domain 2) and a macaque-specific domain (domain 15), which were clearly separated in the embedding space (**Supplementary Fig. 12a, b**). Given that amyloid-*β* (A*β*) plaque accumulation is a hallmark of AD pathology, we examined the relationship between the mouse-specific domain and A*β* deposition. Domain 2 exhibited a strong spatial overlap with regions of high A*β* immunostaining intensity (**Fig. 5c, d**). Quantitative comparison against randomly sampled control spots of equal number confirmed significantly higher A*β* plaque intensity in domain 2 (**Fig. 5c**). Differential expression analysis further revealed that genes enriched in the mouse AD-associated domain, such as *CTSS, CST3*, etc. It showed disease-specific spatial expression patterns that were absent in the macaque hippocampus (**Fig. 5e**; **Supplementary Fig. 12c**), indicating that STACAME can disentangle pathological domains from conserved anatomical structures through cross-species integration.

**Figure 5:**
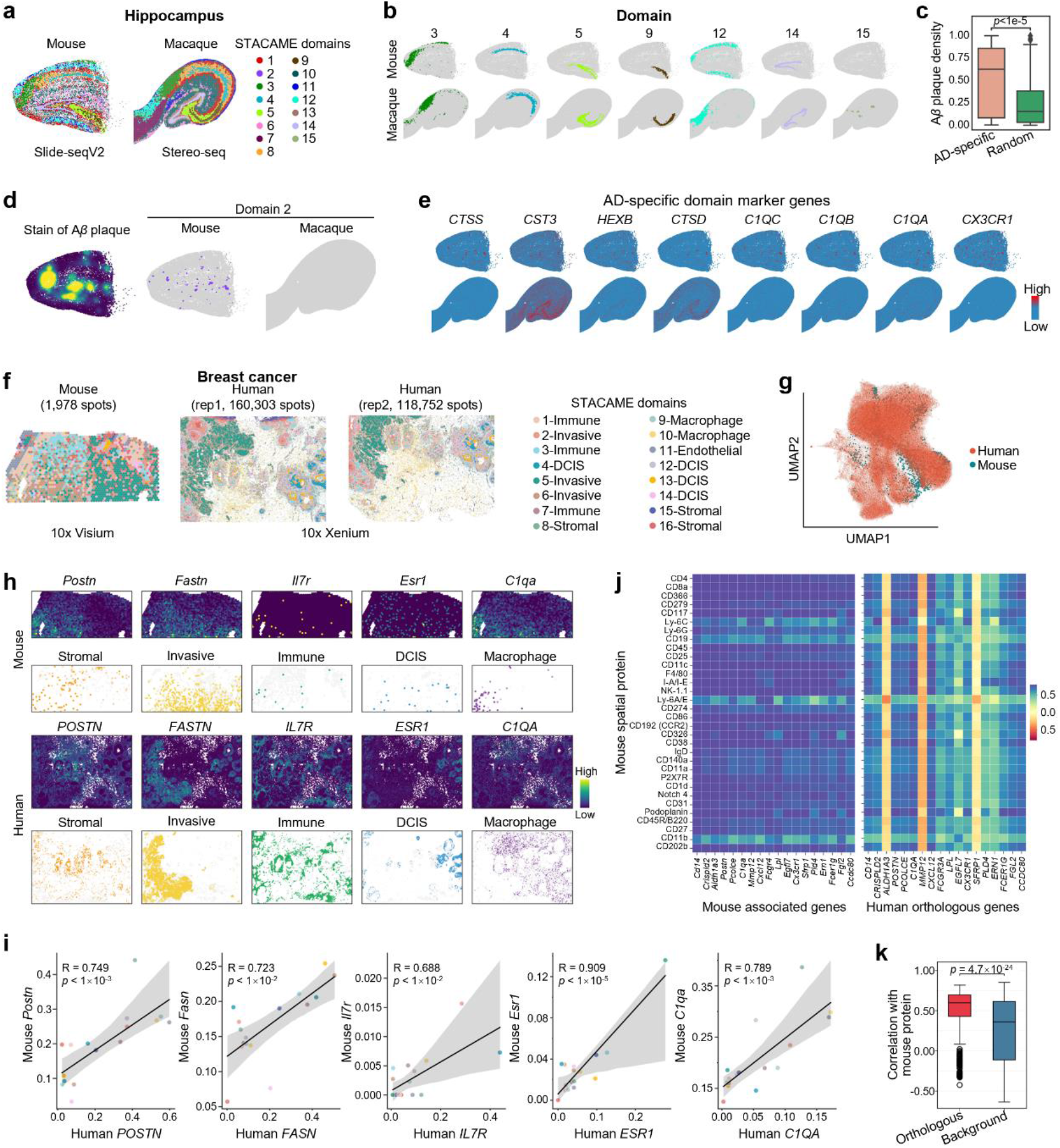
Cross-species integration in disease contexts: Alzheimer’s model and breast cancer. **a–e**, Integration of normal macaque hippocampus (Stereo-seq) and mouse hippocampus (Slide-seqV2) under Alzheimer’s disease (AD) conditions^80^. **a**, Spatial domains assigned by *mclust* on STACAME embeddings. **b**, Species-shared domains (each representing homologous regions) and one macaque-specific domain (domain 15). **c**, Comparison of amyloid-*β* (A*β*) plaque immunostaining intensity in mouse domain 2 versus randomly selected control spots (same number). *P* value is calculated by the Welch’s *t*-test with Bonferroni correction. **d**, Adjacent tissue section showing Aβ plaque distribution (left), spatial map of domain 2 in mouse (middle), and in macaque (right). **e**, Spatial expression patterns of the top differentially expressed genes (DEGs) in mice versus macaques. Using the Wilcoxon rank-sum test with log-transformed expression values, genes were ranked by their differential expression between the target domain and all other domains, with the raw expression matrices used in the calculations. Significantly DEGs were identified using stringent thresholds: false discovery rate (FDR) < 0.01 and log_2_(FC) > 1.5. **f–k**, Integration and analysis of mouse (10x Visium) ^51^ and human breast cancer (BC, 10x Xenium)^52^ tissues. **f**, 16 shared spatial domains identified by STACAME, annotated referring to tumor marker genes and domain cell-type compositions. **g**, UMAP of BC spot embeddings. **h**, Spatial maps of invasive tumor, non-invasive tumor, and TME marker gene expression and STACAME relevant domains. **i**, Scatter plots with linear regression lines and shaded 95% confidence intervals for six human-mouse ortholog pairs (normalized expression, common domains only). Shaded regions are 95% confidence intervals (not standard deviation) of the regression fit. Linear regression with a two-tailed t-test (null hypothesis: slope = 0) yielded Pearson (*R*) and *P* value, indicated in each plot. **j**, Heatmaps of domain-level correlations between mouse spatial proteins and genes. Left, correlation matrix of mouse proteins with the top 300 protein-associated mouse genes (screened via Pearson correlation of domain-averaged expression). Right, correlation matrix of mouse proteins with human orthologs matched from these top 300 mouse genes. **k**, Boxplot showing the significance of cross-species domain correlation conservation. Orthologous group: correlations between mouse proteins and human orthologs; background group: correlations between mouse proteins and randomly selected human background genes (sample size-matched). Statistical significance was assessed by the Mann–Whitney U test (rank-sum test; *P* value is shown).

To investigate whether latent AD-related regions could be detected in the normal macaque hippocampus, we removed AD-associated differentially expressed genes from the mouse dataset and repeated the integration. Under this setting, after running *mclust* on the STACAME embeddings again, the AD-specific domain disappeared, while the remaining domains retained clear correspondence to anatomical hippocampal regions (**Supplementary Fig. 13a, b**). Using MNN mapping in the filtered embedding space, we identified macaque spots most closely aligned with the former mouse AD domain (**Supplementary Fig. 13c**). However, these spots did not exhibit significant differential expression relative to other macaque regions (**Supplementary Fig. 13d**), suggesting that AD-associated transcriptional signatures emerge as disease-induced alterations rather than reflecting pre-existing molecular states in the normal macaque hippocampus.

We further applied STACAME to integrate the mouse and human breast cancer (BC) tissues profiled by 10x Visium^51^ and 10x Xenium^52^, respectively. STACAME identified 16 shared spatial domains across species (**Fig. 5f, g, Supplementary Fig. 15**), including three invasive ductal carcinoma (IDC) domains (domains 2, 5 and 6), four domains that represent noninvasive forms (DCIS) of BC (domains 4, 12, 13, and 14), and 9 other domains that belong to tumor microenvironment, which are annotated according to the marker gene enrichment and cell type proportions (**Supplementary Fig. 16**). The stromal, IDC, immune, DCIS, and macrophage domains show conserved expression of key breast cancer marker genes (*Postn, Fastn, Il7r, Esr1, C1qa*) (**Fig. 5h**), which reflects the biological meanings of STACAME domains. *Postn* marks stromal CAFs^52,53^; *Fastn*, highly expressed in IDC/DCIS, supports tumor lipid metabolism; *Il7r* regulates immune cells in the immune domain; *Esr1* is a core *HR*^+^ (Hormone Receptor-positive) breast cancer marker in DCIS/IDC; *C1qa*, expressed in macrophage/immune domains, participates in immune regulation. Their expression reflects domain functional specialization. In addition, these genes display significantly correlated expression across homologous domains between mice and humans (**Fig. 5i**), indicating STACAME domains present conserved tumor or immune programs between mice and humans.

Given that protein–gene transcriptional relationships are critical for breast cancer research, linking genomic alterations to tumor phenotypes and uncovering disease mechanisms, this study conducted a domain-level cross-species correlation analysis integrating spatial proteomics and transcriptomics data. Specifically, Pearson correlation coefficients were computed between domain-averaged mouse antibody-derived tags (ADTs) expression sequenced by SPOTS^51^ (32 ADTs for TME) and gene expression to identify the top 300 protein-associated mouse genes, followed by mapping to their human orthologs. Correlation matrices were then constructed to compare mouse protein-gene associations within mice and across species (**Fig. 5j**). In general, protein-gene correlation patterns are largely conserved between mice and humans across ADTs, reflecting tightly coupled regulation of core lineage and immune-state markers^54^. These include lymphoid identity markers (CD19, CD45R/B220), myeloid markers (Ly-6C, Ly-6G, CD11b), and stem/progenitor or epithelial markers (Ly-6A/E, CD326). Notably, we observed particularly strong cross-species concordance for Ly-6C, Ly-6G, CD19, Ly-6A/E, and CD11b, supporting evolutionary conservation of myeloid and lymphoid immune programs within corresponding spatial domains. In contrast, while the mouse orthologs of *ALDH1A3, MMP12*, and *SFRP1* show positive correlations with ADT profiles, their human counterparts display consistent negative correlations across all 32 ADTs. These genes are functionally linked to tumor and stromal programs, including cancer stemness (*ALDH1A3*)^55^, macrophage-driven extracellular matrix remodeling (*MMP12*)^56^, and Wnt signaling antagonism and tissue homeostasis (*SFRP1*)^57^, rather than canonical immune cell identity. Such inverse correlations likely reflect spatial segregation between tumor/stromal and immune compartments, divergent post-transcriptional regulation between species, and species-specific wiring of stromal-tumor crosstalk, rather than core immune programs. Furthermore, the conservation of these domain-level correlations was quantitatively evaluated by comparing orthologous gene pairs with randomly sampled background genes using a Mann-Whitney U test, thereby assessing the statistical significance of cross-species transcriptional coupling (**Fig. 5k**).

Consistent patterns of conserved and divergent disease-associated spatial domains were also observed in the cross-species integration of injured mouse liver tissue (Stereo-seq, D17)^58^ and normal human liver tissue (10x Visium) (**Supplementary Fig. 17a, b**). Cholangiocyte-rich domains were identified based on the expression of *Krt19* and *Krt7*, and both the cholangiocyte domain (domains 2, 3) and the liver progenitor-like cell (LPLC) domain (domains 4, 6 and 10) exhibited increased spatial representation in the injured mouse liver, in agreement with prior studies^58,59^. These domains showed elevated activity of *TGFβ* signaling, consistent with the role of cholangiocytes in driving progenitor cell differentiation and modulating hepatocyte proliferation during liver injury and repair. In the mouse liver, STACAME identified expanded cholangiocyte and LPLC domains with broader spatial coverage, with domains 2 and 3 showing substantial overlap with regions of high *Krt19* and *Krt7* expression. By contrast, *Tgfb2* expression was largely restricted to the mouse liver and spatially coincided with the identified domain 12, further highlighting disease-specific molecular signatures captured by the model (**Supplementary Fig. 17c–f**). Notably, STACAME also detected a small human-specific domain enriched for gene programs associated with hepatocellular carcinoma and classical monocytes, suggesting a potential subclinical pathological state in the human liver sample (**Supplementary Fig. 18**).

Together, these results demonstrate that STACAME enables robust identification of disease-associated spatial domains while preserving conserved tissue architecture, facilitating comparative analysis of pathological mechanisms across species and disease contexts.

### STACAME enables large-scale, robust 3D integration of ST data across species

To evaluate the scalability of STACAME for large-scale and 3D cross-species ST data, we applied it to cerebellar datasets from mice, marmosets, and macaques, comprising more than 10 million spatial spots across 98 tissue sections. The embeddings generated from STACAME revealed clear separation of major cerebellar layers across species, as visualized by UMAP colored by four manually annotated layers, i.e., molecular, Purkinje, granular, and white matter (**Fig. 6a**). Clustering of the integrated embeddings using *mclust* showed strong agreement with manual layer annotations, as reflected by high ARI values (**Fig. 6b**, left). Increasing the clustering resolution to k = 7 further subdivided the molecular layer into four spatially distinct subclusters for all species together (**Fig. 6b**, middle and right) and for each species (**Supplementary Fig. 19a**), indicating finer-grained organization within this layer. And the batch effects among the slices within and across species are removed visually (**Supplementary Fig. 19b**). 3D reconstruction of the inferred domains (k=4) by ICP and Spateo^60^ further demonstrated that STACAME robustly captures shared cerebellar architecture despite substantial interspecies differences in size and morphology (**Fig. 6c**; **Supplementary Fig. 19c,d**). Analysis of normalized domain composition across species identified three species-specific domains (domains d2, d3, and d5), whose normalized domain composition value is close to or larger than 0.5, whereas the remaining domains were conserved across all three species (**Fig. 6d**; **Supplementary Fig. 20a, b**). The slice specificity of individual domains is also revealed (**Supplementary Fig. 20c**), which is consistent with the variance of cerebellar slices.

**Figure 6:**
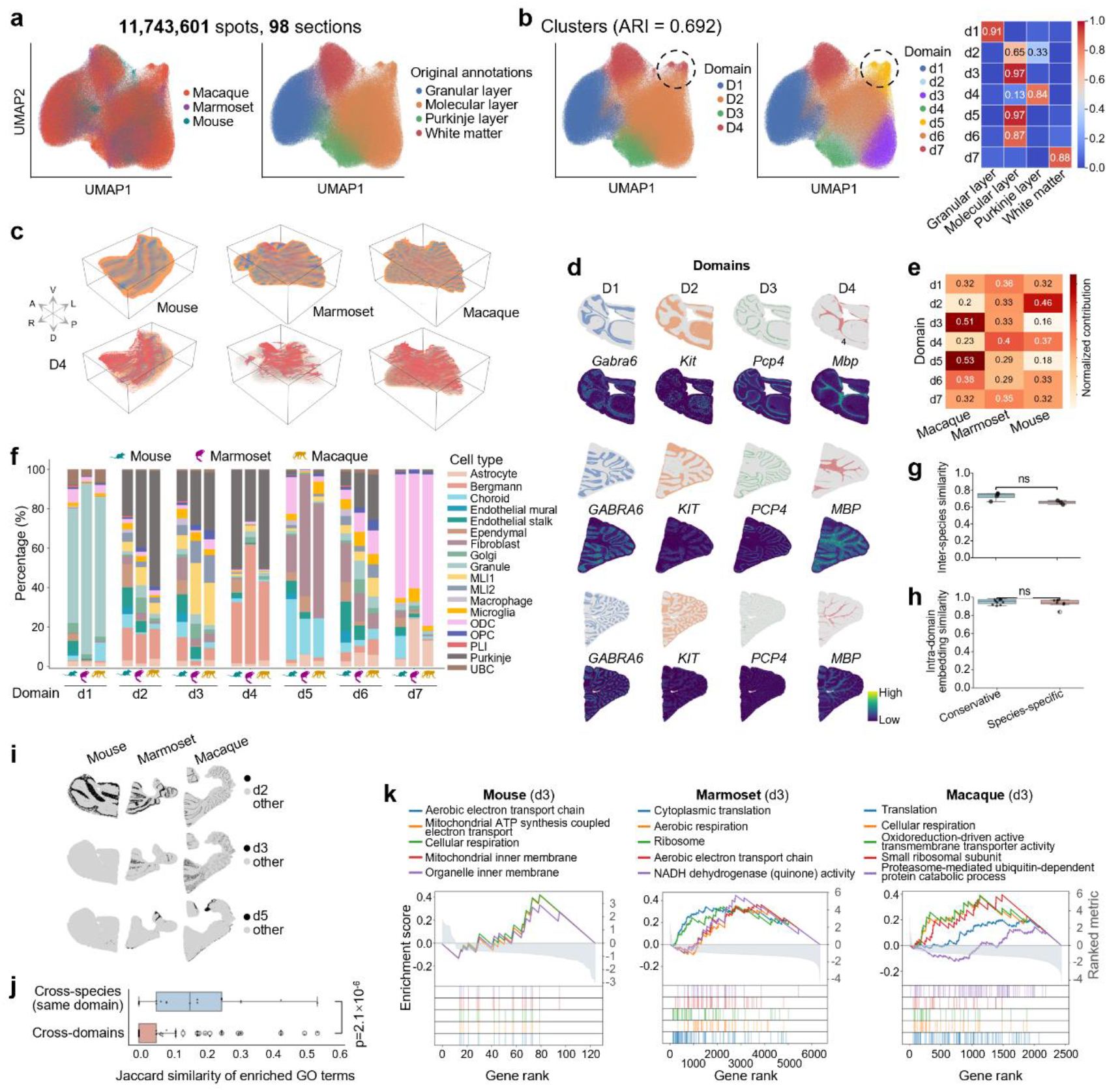
Large-scale cross-species 3D integration of cerebellar spatial transcriptomics (>10 million spots). **a**, UMAP of spot embeddings colored by four annotated cerebellar layers. **b**, UMAP of domains with ARI (left) indicating high concordance with manual annotations (except a minor cluster, dashed circle). When clustering resolution *k* = 7, the molecular layer splits into four subclusters (middle), as indicated in the confusion matrix (right). **c**, 3D reconstruction of common domains identified from STACAME embeddings (top: four domains; bottom: D4 corresponding to white matter). **d**, Normalized domain composition across species reveals three species-specific domains (2, 3, and 5). **e**, Spatial maps of four domains matching annotated layers and their canonical marker genes. **f**, Cell-type composition across the seven domains (boxplots as in **Figure 5**). **g**, Inter-species similarity of cell-type proportions is higher in conserved domains (1, 4, 6, 7) than in species-specific ones (2, 3, 5). **h**, Intra-domain embedding similarity is also greater in conserved domains. **i**, Spatial localization of species-specific domains (T175 in mouse, T490 in marmoset, T62 in macaque). **j**, Jaccard similarity of the top 10 enriched GO terms. Within-domain comparisons (*n* = 21) show significantly higher similarity than between-domain pairs (*n* = 189; one-sided Mann-Whitney *U* test, *P* =2.14×10^−6^). For each domain and species, domain-specific DEGs were identified using the Wilcoxon rank-sum test (*P*<10^−2^) and ranked by log_2_(FC). Pre-ranked gene set enrichment analysis (GSEA) was performed against human biological process, cellular component, and molecular function gene sets with 1000 permutations. Only terms with positive normalized enrichment scores (NES > 0) and FDR *q*-values < 0.05 were retained, and the top 10 terms ranked by FDR were used for downstream similarity analysis. **k**, Gene rank plots for upregulated GO terms in domain 3, demonstrating high cross-species functional conservation. Marker genes were ranked by log_2_(FC) from a Wilcoxon rank-sum test. Preranked GSEA (GSEApy, 1,000 permutations) was conducted against the GO Biological Process 2023, GO Cellular Component 2023, GO Molecular Function 2023, Allen Brain Atlas up, Azimuth Cell Types 2021, and CellMarker Augmented 2021 libraries. Significantly upregulated gene sets (FDR q‐value < 0.05, NES > 0) are plotted; the top GO terms are shown.

Spatial maps of representative shared domains demonstrated consistent anatomical localization and expression of canonical marker genes across species (**Fig. 6e**). For each domain (k=7), the cell type compositions are visually consistent, with only minor variations in the proportions of rare cell types, even in the species-specific domains (**Fig. 6f**). The divergence in cell-type composition is greater between mouse and the primate species than between marmoset and macaque, which is consistent with their phylogenetic distances and evolutionary relationships, and further supports the biological validity of the spatial domains identified by STACAME. Further cell-type composition analysis revealed that conserved domains (domains d1, d4, d6, and d7) exhibited no significantly higher inter-species similarity in cell-type proportions than species-specific domains (i.e., domains d2, d3, and d5) (**Fig. 6g**). Consistently, intra-domain embedding similarity was also not significantly higher in conserved domains (**Fig. 6h**). These results indicate stronger spatial and molecular coherence between species-specific domains and species-shared domains, and the STACAME domains are consistently valid. Sankey and enrichment analyses further highlighted preservation of cellular architecture within conserved spatial niches across evolution (**Supplementary Fig. 21**).

Species-specific domains displayed distinct spatial localization patterns within each cerebellum (mouse T175, marmoset T490, and macaque T62; **Fig. 6i**; **Supplementary Fig. 22**), reflecting evolutionary divergence. The Jaccard similarity of top 10 enriched Go terms ranked by FDR demonstrated significantly greater similarity of top enriched Gene Ontology (GO) terms within the same domain across species compared to between-domain pairs (one-sided Mann–Whitney U test, *P* =2.14 × 10^−6^; **Fig. 6j**). Gene rank analyses of upregulated GO terms in representative domains, including domains d2, d3, and d5, further confirmed strong cross-species conservation of functional programs despite anatomical divergence (**Fig. 6k**; **Supplementary Fig. 23**). For instance, cellular respiration and aerobic electron transport chain are conserved across species for domain d3, indicating its cellular function.

Together, these results demonstrate that STACAME scales efficiently to tens of millions of spatial observations and enables robust three-dimensional integration of spatial transcriptomics across species, preserving conserved cerebellar organization while resolving species-specific spatial and functional specializations.

### STACAME provides a reliable framework for cross-species cell-type discovery and annotation

A key application of cross-species spatial transcriptomic integration is the prediction of cell types in one species based on annotations from another. To evaluate this, we applied STACAME to integrate macaque dorsolateral prefrontal cortex (DLPFC, Stereo-seq)^37^ and human prefrontal cortex (PFC, Slide-tags)^61^, both with manually annotated cortical cell types (**Supplementary Fig. 24a**). After obtaining the STACAME embeddings, a two-layer fully connected neural network (**Methods**) was trained on macaque embeddings to predict cell types in human embeddings, and vice versa. Experiments were repeated five times with different random seeds to account for stochastic variability. For comparison, we applied several baseline integration methods, while the supervised STELLAR model^62^ directly predicted human cell types using macaque embeddings.

STACAME consistently outperformed all baselines, achieving the highest prediction accuracy and macro F1 score in both directions (macaque-to-human and human-to-macaque; **Supplementary Figs. 24b, c and 25a, b**), with statistically significant improvements over STELLAR (two-sided Welch’s t-test with Bonferroni correction, P < 0.001). Predicted cell types were spatially coherent and matched known marker gene expression patterns (**Supplementary Fig. 24d**), and embedding correlations between homologous cell types were substantially higher than non-homologous pairs (**Supplementary Fig. 24e**).

We further extended this analysis to cerebellar slices from mice, marmosets, and macaques (**Supplementary Fig. 25f**). Cross-species prediction using STACAME embeddings from the other two species achieved high accuracy and macro F1 scores, demonstrating robust transferability across both cortical and cerebellar regions (**Supplementary Fig. 24f, g**). Embedding correlations between homologous cell types were substantially higher than non-homologous pairs (**Supplementary Fig. 25c**). Additional analyses confirmed that while increasing the amount of spatial neighbor information can slightly reduce standard metrics (accuracy, F1, precision, recall), STACAME embeddings remain highly informative for cell-type prediction (**Supplementary Fig. 25d, e**). These results indicate that STACAME provides a reliable framework for cross-species cell-type annotation for ST data.

### Ablation experiments validate the design choices of STACAME

We conducted systematic ablation studies to investigate the contributions of the key components in STACAME and validate the design decisions. First, we examined the impact of including non-homologous genes on integration. When integrating the macaque and human DLPFC datasets, integration performance metrics remained stable with less decay than the other methods, such as STAligner, as the proportion of non-homologous genes increased (**Supplementary Fig. 26a**), demonstrating that STACAME could robustly handle species-specific genes without compromising integration quality.

We further assessed how non-homologous genes influence latent representations by training a two-layer fully-connected neural network to reconstruct original gene expression from embeddings. Using mouse and macaque hippocampus data with the training to test ratio = 8:2 (10 independent repeats), STACAME achieved significantly lower reconstruction loss compared to alternative methods (Welch’s t-test with Bonferroni correction; **Supplementary Fig. 26b**). This indicates that STACAME effectively preserves species-specific biological information within the joint embedding space rather than discarding it during integration.

To evaluate the importance of many-to-many homologous relationships in distantly related species, we analyzed the mouse (E11.5) and zebrafish (24hpf) embryonic datasets (**Supplementary Fig. 26c**). Notably, the number of many-to-many homologous genes (excluding one-to-one orthologs) was comparable to that of one-to-one orthologs, highlighting their quantitative significance. We then compared integration performance using only one-to-one orthologs versus both one-to-one and many-to-many orthologs for constructing cross-species links (20 replicates with different random seeds). Integration with the expanded ortholog set significantly improved species mixing scores (*P* = 8.031 × 10^−5^; **Supplementary Fig. 26d**), confirming that incorporating diverse homology types enhances spatial alignment accuracy between evolutionarily distant organisms.

Finally, we systematically evaluated the objective function components of STACAME and preprocessing steps (**Supplementary Fig. 27a**). Ablation of individual loss terms (i.e., multi-scale discrepancy (distribution alignment), adversarial (domain confusion), and triplet (intra-species discriminability)) revealed their complementary roles, and each contributes uniquely to integration quality and biological structure preservation. Similarly, removing key preprocessing steps (highly variable gene selection, *combat* batch correction, PCA projection before mutual nearest neighbor alignment, or concatenated input construction) consistently degraded performance metrics. The radar plots summarizing preprocessing steps and loss-term ablations illustrate how balanced weighting of all three components achieves optimal trade-offs between integration metrics and biological fidelity (**Supplementary Fig. 27b**). These comprehensive analyses validate STACAME’s architectural design and highlight its robustness across diverse experimental configurations.

## Discussion

In this work, we present a three-step, self-supervised framework, STACAME, designed to overcome the pronounced heterogeneity inherent in cross‐species spatial transcriptomics datasets. We demonstrate the versatility and robustness of STACAME through comprehensive analyses of 13 ST datasets spanning distinct tissues, developmental stages, and disease models across humans, macaques, marmosets, mice, and zebrafish, generated on diverse experimental platforms. Moreover, STACAME exhibits scalability, handling over ten million cells in a cerebellar integration experiment.

To our knowledge, STACAME represents the first computational approach specifically tailored for multi‐species ST integration that simultaneously delineates shared and species‐specific spatial domains. Our findings thus align with and extend recent efforts in comparative cross‐species genomics, underscoring how integrated spatial frameworks can reveal fundamental principles of tissue organization shared across species, while also pinpointing key transcriptional and architectural divergences. This capability not only advances cross-species studies but also provides a translational bridge for leveraging model organisms to interpret human biology and disease.

The proposed CDSVGs and CCSVGs are natural consequences of spatial genomic divergence among species (**Fig. 3**). The procedure does not account much for domain relations and those homologous genes that are not differentially expressed on certain domains. Besides, it may be sensitive to the DEGs or SVGs identification algorithm. Moreover, a proper way is supposed to identify the CDSVGs in the process of cross-species ST integration, since those genes may bring noise for shared domain identification. Hence, iteratively pruning the gene orthologs network according to the CDSVGs could be an effective way to improve the domain alignment, even though STACAME already achieves it by inducing species-specific genes. In addition, the spatially divergent homologous genes probably affect the identification of cell-cell communication65 or gene regulatory networks66. Thus, the cross-species ST integration provides an opportunity to study the cause of differences among species tissues. As for the other omics, such as histological images and spatial epigenomic data, the cross-species integration is also available, but requires more designs for STACAME.

## Methods

### Data preprocessing

STACAME first processes the spatial transcriptomics data from different species with multiple ST slices per species. For each species, we identified the set of genes common to all slices to ensure consistency and concatenated the corresponding raw gene expression matrices. We then retrieved the many-to-many gene orthologs across species according to the gene orthologs database, which were subsequently used to build the cross-species spot adjacency graph. The raw gene expressions were normalized based on library size and log-transformed using the SCANPY package^63^.

#### Gene selection

Highly variable genes (HVGs) were selected for downstream procedures. There are two separate gene selection settings for species links and feature representation. In species link setting, we extracted all the many-to-many orthologous genes across all species and concatenated the many-to-many gene matrices for all the species (*X*_*e*_), and notably, there might be replications of genes to handle many-to-many genes (For example, one gene repeats three times to match its three various orthologs). Then the top HVGs are also selected in this gene matrix to filter out noise. In the feature representation setting, we selected a part of the top highly variable genes (default 1000) for each species, which includes many-to-many gene orthologs, combined with the top highly variable one-to-one homologous genes. For the one-to-one genes across all species, we selected the top 3,000 to 6000 highly variable genes for different tasks. The whole gene expression feature vector is the direct concatenation of the two parts. The SCANPY built-in function *highly_variable_genes()* was used to identify the top genes with the highest dispersions as HVGs. The input of the auxiliary lightweight graph attention autoencoder is PCA-transformed top highly variable many-to-many orthologous gene expression data *X*_*e*_, which contains repeats of genes for aligning the feature dimension.

#### Gene orthology

We downloaded the gene many-to-many homology (orthologs) information from the *BioMart* web server derived from the *Ensembl Compara* pipeline^64^ following the previous work^10^. We used the mouse as the anchor species and downloaded the homology mapping file.

### Construction of the spatial neighbor graph

For each slice, two spots are considered as neighbors if the Euclidean distance between their spatial coordinates is below a pre-defined threshold *r*. We construct an undirected neighbor graph denoted as the adjacency matrix ***A***, where ***A***_*ij*_ = 1 if and only if spot *i* and spot *j* are mutual neighbors. ***A*** is a symmetric matrix with self-loops added for each spot. The value of *r* can be adjusted so that each spot has 3–10 nearest neighbors on average for diverse ST scenarios and platforms. We use *G*_*p*_ to denote the concatenated neighbor graph across species.

### Construction of cross-species spot adjacency graph

To establish adjacent edges among spots from different species, we first retrieved all the many-to-many homologous genes referring to the downloaded gene orthologs database. Then, we systematically traverse all possible gene combinations with each member originating from a distinct species. By extracting and concatenating expression values corresponding to these enumerated combinations for every spot, we obtain length-aligned feature vectors that enable consistent cross-species comparison. We implement MNN or KNN (optional) to build cross-species edges. For MNN, we apply three optional steps before computing the Euclidean distance matrix across species: HVGs selection with *highly_variable_genes()* in SCANPY, batch effect removal with *sc*.*pp*.*combat()*, and dimension reduction with PCA. MNN is executed on the processed matrix, and a number of cross-species mutual pairs are obtained. For KNN edges, each spot’s *K* nearest cross-species neighbors are selected via cosine similarity directly.

Notably, the final spot neighbor graph is composed of two types of edges: intra-species spatial edges from spatial nearest neighbors within each slice, and cross-species expression edges derived from MNN (or KNN) analysis. The KNN edges are optional and practically useful when the heterogeneity among species is extremely high.

### Graph attention auto-encoder for pretraining

The model training process consists of two stages: the pretraining stage and the species alignment stage. In the pretraining stage, the combined features, including one-to-one homologous genes and species-specific gene expression values, are first fed into a graph attention autoencoder to get the initial lower-dimensional embeddings of the spots. In the species alignment stage, the pretrained embeddings for all the species are concatenated as the input of a specially designed auto-encoder whose encoder is trained to aligning the pretrained embeddings.

Suppose there are *S* species in the dataset, and species *s* contains *N*_*s*_ spots, *s* = 1, 2, …, S. Let *L*_*p*_ be the number of layers in the encoder in the pretraining stage. For species *s*, the output representation of the *k*th (*k* ∈ {1,2, …, *L*_*p*_ − 1}) encoder layer of spot *i* ∈ {1,2, …, *N*_*s*_} is formulated as follows:

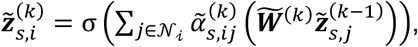

where *N*_*i*_ represents the neighbor set of spot *i* (including spot *i* itself), 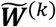 is the trainable weight matrix, *σ* is the nonlinear activation function, 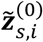 is the initial gene expression ***x***_*s,i*_ of spot *i* in species *s*. 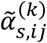 is set to normalize the relevance coefficients of the neighbors of spot *i*, which are also called graph attention coefficients:

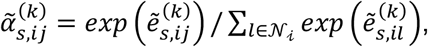

where 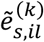 represents the relevance of spot *i* to its neighboring spot *j* of the species *s*, and is computed by:

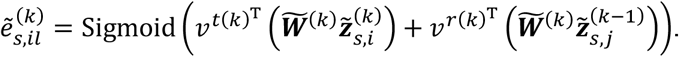

Here, Sigmoid denotes the sigmoid activation function, and ***v***^*t*(*k*)^ and ***v***^*r*(*k*)^ are the trainable weight parameters of the *k* th layer. The *L*_*p*_-th encoder layer does not use the attention mechanism and is calculated by:

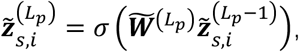

which is considered as the pretrained embedding of spot *i* in the dataset of species *s*.

The layer number of the decoder is the same as that of the encoder. The decoder reverses the process of the encoder to reconstruct expression features. The decoder uses the output of the encoder as the input, and the *k*th (*k* ∈ {2, …, *L*_*p*_ − 1, *L*_*p*_}) decoder layer reconstructs the representation of spot *i* in layer *k* − 1 as 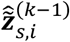. Similar to the encoder, the attention mechanism is not used in the last layer. The output 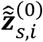 is considered the reconstructed expressions. For each species *s*, the reconstruction loss during the pretraining stage is

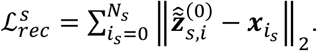

### Graph attention auto-encoder for alignment

In the species alignment stage, the pretrained embeddings of all *S* species are collected and concatenated to a set of a total number of *N* spots, i.e.,

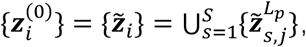

where *s* = 1,2, …, *S*,*j* = 1,2, …, *N*_*s*_ and *i* = 1,2, …, *N*. The initial gene expression of spot *j* in species *s*, i.e., {***x***_*s,j*_|*s* = 1,2, …, *S, j* = 1,2, …, *N*_*s*_}, are also concatenated to {***x***_*i*_|*i* = 1,2, …, *N*}. Then let *L*_*a*_ be the number of layers in the encoder and *N* be the number of spots in the slices.

Different from the pretraining GAE, the input dimension of the aligning GAE is the latent dimension of the pretraining GAE, and the output is the reconstructed expression data, which could also be PCA of the gene expression matrix for computational efficiency. Moreover, since the asymmetric architecture, the decoder does not reverse the process of learning latent embedding in the encoder to reconstruct expression features. Suppose the decoder output 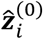 is the reconstructed gene expression data, the reconstruction loss of the aligning stage is

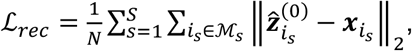

where *M*_*s*_ is the spot index set of species *s*.

The merged expression matrix used for the alignment stage consists of a common block of highly variable homologous genes, followed by contiguous species‐specific blocks. Each species‐specific block was zero‐padded to match the maximum number of species‐specific genes across all species, so that for every spot, only its own species’ block contained non‐zero expression values while the remaining blocks were zero. This construction ensures that the feature dimensions of all spots are consistent, and to improve computational efficiency, the resulting matrix can be (optionally) reduced through PCA.

### Cross-species triplets learning

To remove the data heterogeneity across species, we employed cross-species triplet learning, which has shown superior performance in diverse fields, including ST data integration^27,65^. Since the induced species-specific genes bring disturbance for computing MNN, we utilize an auxiliary lightweight graph attention autoencoder using the PCA of the expression matrix (*X*_*e*_) generated by mapping orthologous genes across species. Cross-species triplets are constructed by identifying inter-species MNN spot pairs in both the gene expression space and the latent space.

To leverage the constructed triplets for model training, the anchor–positive pairs are used to overcome the batch effect while the anchor–negative pairs guide the alignment in a discriminative way. The anchor-positive pairs are composed of MNN (KNN is optional) spots that have similar gene expression from two different species, while the anchor-negative pairs are randomly selected from the dataset of the same species. The cross-species triplet loss is defined as:

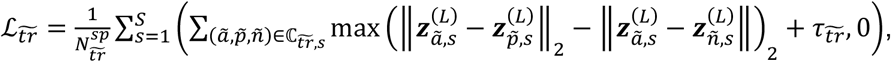

where 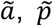, and ñ represent the indices of the anchor, positive and negative spots respectively, 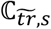 is the set of the identified spot triplets with size 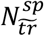 and 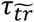 (default 1.0) is the margin used to enforce the distance between positive and negative pairs.

For situations where some species datasets contain multiple sections, we also implement a cross-slice triplet loss as proposed in STAligner^27^. The MNN pairs are used to define the anchors and positive spots that have similar gene expression but are from two different slices, and negative spots are randomly selected from the same sections. The cross-slice triplet loss is defined as:

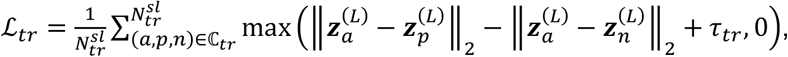

where *a, p*, and *n* denote the indices of the anchor, positive, and negative spots, respectively, ℂ_*tr*_ represents the set of identified spot triplets with a cardinality of 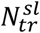, and *τ*_*tr*_ (default value 1.0) serves as the margin utilized to enforce the separation distance between positive and negative pairs.

### Latent space alignment with domain adaptation loss

Maximum Mean Discrepancy (MMD) loss is a crucial component in unsupervised domain adaptation^66,67^, employed to minimize the distributional divergence between the source and target domains. By incorporating MMD loss into the learning objective, STACAME learns domain-invariant features among species, effectively reducing the distance among domain feature distributions of multiple species, thereby improving the model’s ability to transfer knowledge from one species’ domain to the domains of other species. Specifically, the MMD loss in STACAME is defined as:

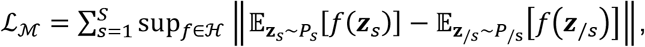

where *f* denotes a function in Hilbert space ℋ, *P*_*s*_ represents the distribution of embedding ***z***_*s*_ of species *s*, and *P*_/*s*_ represents the distribution of embedding ***z***_/*s*_ of the other species. We subsample a batch of data from each species for MMD loss, for computational efficiency.

To complement the kernel-based distribution alignment imposed by MMD loss, we additionally incorporate an adversarial learning objective as an optional loss to further reduce residual cross-species discrepancies in the latent space. As MMD may be less sensitive to complex, higher-order differences when species exhibit pronounced transcriptional or spatial heterogeneity, adversarial learning provides a flexible, data-adaptive mechanism that addresses this limitation by aligning latent distributions in a discriminative manner. A species discriminator is trained to predict the species identity of latent embeddings. In contrast to MMD, which enforces alignment through explicit moment matching, the adversarial loss implicitly captures higher-order distributional differences by optimizing against a learned decision boundary. As a result, the two objectives operate at complementary levels: MMD promotes global statistical consistency across species, whereas the adversarial loss refines local and higher-order alignment by penalizing species-specific separability in the embedding space. Formally, let *D*_*s*_ be the species discriminator. Let ***z*** represent the latent embeddings from all other species. During optimization, the discriminator is trained to maximize

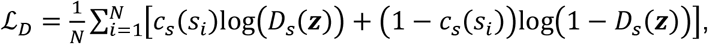

while the GAE of the aligning stage is trained adverbially to minimize it, thereby encouraging species-invariant embeddings. The adversarial loss is defined as the

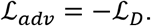

This complementary design for jointly optimizing MMD and adversarial objectives improves the stability of training and the model’s ability to capture both conserved and divergent biological signals, ultimately leading to a more coherent and transferable shared embedding space across species.

### Manifold preserving loss

To prevent the cross-species alignment from distorting the intrinsic local topology of each species, we introduce a manifold-preserving loss that enforces consistency between the original pretraining embedding space and the aligned latent space. We randomly sample a set of intra‐species edges *E* at each epoch and impose a weighted Laplacian penalty. For each edge (*i, j*), a Gaussian affinity 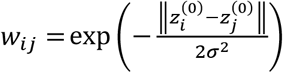 is first computed from the pretrained embeddings, with *σ*^2^ set to the median squared distance over *E*. To prevent larger species from dominating the loss, each edge is further multiplied by a balancing factor 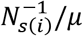, where 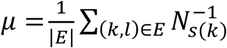 is the mean reciprocal species size over the sampled edges. The loss is applied symmetrically to both the main embedding ***z*** and the auxiliary embedding ***z***^aux^ as

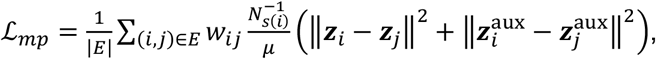

and its overall contribution is weighted by a non‐negative hyperparameter *γ*_mp_.

### The STACAME loss

The overall loss function of STACAME is the weighted sum of the reconstruction loss, cross-slice triplet loss, cross-species triplet loss, MMD loss and GAN loss (optional) as follows:

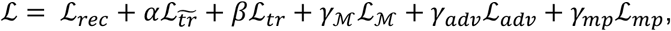

where *α, β, γ*_*M*_,*γ*_*adv*_ and *γ*_*mp*_ are non-negative real numbers with default values of 1 and *β* is set to 0 when each species contains one single slice.

### Benchmarking methods

We considered the following integration methods to benchmark the integration performance: Harmony^42^, Scanorama^68^, SEDR^43^, and STAligner^27^. We also used SCANPY^63^ to produce uncorrected embeddings of the combined raw input data, where principal component analysis (PCA) was employed for dimensionality reduction and no integration method was used. For spatial cell type prediction benchmarking, we adopted STELLAR^62^, a supervised cell type prediction method for single-cell ST data using a graph convolutional neural network. All parameters were optimized according to the datasets (Supplementary material). Notably, for all these algorithms, a SCANPY combat batch correction step is supposed to execute in advance. Moreover, only one-to-one gene orthologs are used in the benchmarking methods because of their model mechanisms.

### Evaluation metrics on integration

An effective integration method should accurately identify tissue structures of each species and align corresponding homologous spatial domains across species. Therefore, its resulting spot embeddings should preserve biological variation while simultaneously ensuring thorough mixing of species. Inspired by previous research^69^, we defined biological conservation and species mixing metrics for assessing cross-species data integration, with each metric consisting of multiple sub-metrics. Together, these collectively form an overall integration score for global evaluation.

*Mean average precision (MAP)* was used to evaluate the spatial domain resolution. Suppose that the domain of the *i* th spot is *s*^(*i*)^ and that the spatial domain of its *K* ordered nearest neighbors are 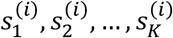, the mean average precision is then defined as follows:

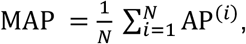

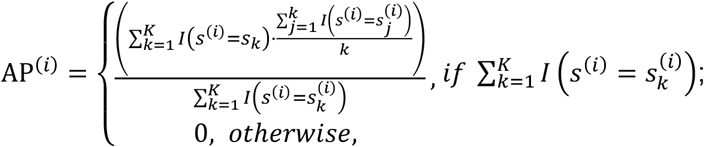

where 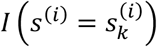 is an indicator function that equals 1 if 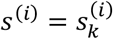, and 0 otherwise. For each spot, average precision (AP) computes the average spatial domain precision up to each domain-matched neighbor, and mean average precision is the average precision across all spots. We set *K* to 1% of the total number of spots in each dataset. Mean average precision has a range of 0 to 1, and higher values indicate better domain resolution.

*Spatial domain ASW (average silhouette width)* was also used to evaluate the domain resolution^70^:

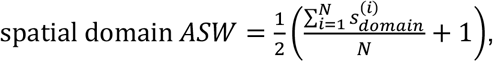

where spatial domain *ASW* is the spatial domain silhouette width for the *i*th spot, and *N* is the total number of spots. The spatial domain ASW has a range of 0 to 1, and higher values indicate better spatial domain resolution.

*Spatial neighbor consistency (SNC)* was used to evaluate the preservation of single-species data variation after multi-species integration and was defined following a previous study^71^:

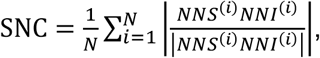

where *NNS*^(*i*)^ is the set of spatially *k*-nearest neighbors for spot *i* in the single-species data, *NNI*^(*i*)^ is the set of *K*-nearest neighbors for the *i* th spot in the integrated space, and *N* is the total number of spots. We set *K* to 1% of the total number of spots in each dataset. The spatial neighbor consistency has a range of 0 to 1, and higher values indicate better preservation of data variation.

#### Biology conservation

The mean average precision, spatial domain ASW and neighbor consistency all measure the biology conservation of data integration. Following the procedure from the recent benchmark study^70^, we compute the average across the three to summarize them into a single metric representing biology conservation:

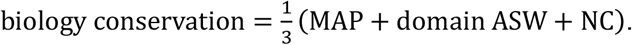

#### Seurat alignment score (SAS)

We adopted SAS to evaluate the alignment of spots or genes in the embedded space, that is, the mixing of embeddings between species. SAS ranges from 0 to 1, and higher values indicate better mixing. Here, we calculated SAS as described in the reference^72^ with the Python implementation^73^. SAS is defined as:

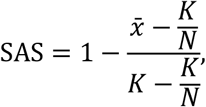

where 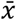 is the average number of spots from the same species among the K-nearest neighbors (different datasets were first sub-sampled to the same number of spots as the smallest data set), and *N* is the number of species. We set *K* to 1% of the sub-sampled spot number. The average SAS is defined as the average of SAS on each homologous domain or region pairs (groups) among species.

*Graph connectivity (GC)* was also used to evaluate the extent of mixing among omics layers and was defined as in a recent benchmark study^70^:

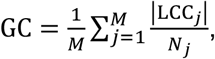

where LCC_*j*_ is the number of cells in the largest connected component of the spot k-nearest neighbors’ graph (K=15) for the spatial domain *j, N*_*j*_ is the number of spots in the spatial domain *j* and M is the total number of spatial domains. The graph connectivity has a range of 0 to 1, and higher values indicate better mixing.

#### Species mixing

Seurat alignment score and graph connectivity all measure species mixing in the data integration. Following the procedure from the recent benchmark study^70^, we first conduct min-max scaling for each of the metrics, and then compute the average across the three to summarize them into a single metric representing species mixing:

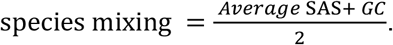

#### Overall integration score

To compute an overall integration score, we use a 6:4 weight between biology conservation and species mixing, following the recent benchmark study:

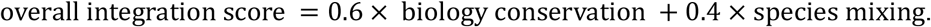

### The training settings of STACAME

In the experiments, the embedding dimension is set to 32 to 64 (default 32). We used the Adam optimizer^74^. For each experiment, hyper-parameters such as learning rate (default 0.001), epoch number (default 3000), MNN hyperparameter (default 8), and data normalization settings are optimized and presented in the code.

### Analysis methods

#### Clustering spot embeddings

Generally, we used the *mclust* algorithm^46^ for clustering on integrated cross-species embeddings, acquiring common domains for multiple species.

#### Identifying differentially expressed genes

For genome analysis with respect to DEGs, we used the Wilcoxon test implemented in the package SCANPY^63^ to identify differentially expressed genes for each spatial domain with the FDR threshold of 1% (Benjamin-Hochberg adjustment).

#### Identifying cross-species divergent and shared spatially variable genes

In our work, for one shared domain across two species, we simply rank the DEGs for each species and select those genes that are ranked top in one species and rank last for another species as CDSVGs. CCSGVs are selected by computing the intersection set of DEGs.

#### Classification and reconstruction neural network

Both the domain or cell type classification and gene expression reconstruction neural networks are two-layer fully connected. The input dimension equals the embedding dimension for each algorithm. The hidden layer dimension is 256 and uses *ReLU* as the activation function. For domain or cell type classification, the output dimension depends on the number of classes. For the gene expression reconstruction task, the output dimension depends on the number of genes used by the method.

### Data description

Source data for Figures 2-6 are available with this manuscript. The datasets analyzed in this study are all from publicly available datasets (**Supplementary Table 1**). The Stereo-seq macaque DLPFC dataset is a part of slice T127 of macaque 1 in the whole cortex dataset^37^ and contains 18,375 spots. The manually annotated labels in the original study include six neocortical layers. The 10x Visium human DLPFC dataset was collected from three independent adult samples. We used slice 15,1673 in the experiment. The manually annotated labels include white matter and six neocortical layers^22^.

We downloaded the hippocampus data profiled by Stereo-seq from the Brain Science Data Center at the Chinese Academy of Sciences and selected slices T315 (4,540 spots) and T36 (28,499 spots) as the mouse and macaque data, respectively^75^. We cropped the slice ‘*Puck_200115_08*’ in the mouse hippocampus dataset^20^ sequenced by Slide-seqV2 to 21,919 spots.

The mouse visual cortex (VIS) dataset sequenced by MERFISH consists of 5,995 spots and 234 genes, and the human auditory cortex (AUD) dataset sequenced by MERFISH contains 3970 spots and 4000 genes^76^.

The human embryo slice was profiled by the 10x Visium platform, containing 2,909 spots^77^. The early mouse embryo dataset profiled by Stereo-seq consists of eight stages (E9.5, E10.5, E11.5, E12.5, E13.5, E14.5, E15.5, and E16.5), while only E9.5, E11.5, and E12.5 were used in this paper and contain 5,913, 30,124 and 51,365 spots^21^, respectively. The early zebrafish dataset profiled by Stereo-seq consists of six stages (3.3hpf, 5.25hpf, 10hpf, 12hpf, 18hpf, 24hpf), and the last three stages were used in this work and contain 2,081, 3,048 and 5,271 spots, respectively^78^.

The ST data of normal and AD disease mouse hippocampus slices, both profiled by Slide-seqV2, contain 19,285 and 15,092 spots^20^, respectively. For the AD slice, an adjacent A*β* plaque staining image of the same brain tissue was also collected.

The mouse breast cancer slice^51^ sequenced with 10x Visium contains 1,978 spots for RNA-seq, along with 32 ADTs, and the human breast cancer slice^52^ sequenced with 10x Xenium contains 118,752 spots (sample 1, replicate 2).

The mouse liver slice^58^ with cholestatic injury profiled by Stereo-seq is the D17_FS3 slice with 27,915 spots, and the normal human liver slice^79^ profiled by 10x Visium contains 2265 spots.

In this work, the mouse, marmoset and macaque coronal ST sections^38^ of cerebellum contain 1,810,619 (32 sections), 1,916,578 (32 sections) and 8,016,404 spots (34 sections), respectively.

The human PFC ST section profiled by Slide-tags^61^ contains 12,574 spots. The spatial single-cell data set of mouse, marmoset, and macaque profiled by Stereo-seq is annotated by previous work^38^ and contains 35,753 (T170), 27,390 (T484) and 53,264 (T44) spots.

## Data availability

The datasets analyzed in this study are all obtained from publicly available datasets. The macaque slice T127 covering the DLPFC dataset is downloaded from the link: https://www.braindatacenter.cn/datacenter/web/#/dataSet/details?id=1663381185152036865. The human DLPFC dataset can be accessed in the spatialLIBD package (http://spatial.libd.org/spatialLIBD). The mouse, marmoset, and macaque hippocampus datasets are obtained from the Brain Science Data Center at the Chinese Academy of Sciences (https://braindatacenter.cn/) and are available at this link: https://cstr.cn/33145.11.BSDC.1684593483.1659922723465732098. The mouse hippocampus dataset (ID: CP815) profiled by Slide-seqV2 is downloaded from https://singlecell.broadinstitute.org/. The human embryo data can be accessed at https://heoa.shinyapps.io/code/. The mouse embryo data can be accessed at https://db.cngb.org/stomics/mosta/. The zebrafish embryo data can be accessed at https://db.cngb.org/stomics/zesta/. The normal and Alzheimer’s disease mouse hippocampus data can be accessed at https://singlecell.broadinstitute.org/single_cell/study/SCP815 and https://singlecell.broadinstitute.org/single_cell/study/SCP1663, respectively. The mouse and human breast cancer tissue can be accessed at https://www.ncbi.nlm.nih.gov/geo/query/acc.cgi?acc=GSE198353 and https://www.10xgenomics.com/products/xenium-in-situ/preview-dataset-human-breast. The mouse liver of cholestatic injury is available at https://db.cngb.org/stomics/datasets/STDS0000239, and the normal human liver ST slice can be accessed at https://db.cngb.org/stomics/datasets/STDS0000103. The mouse, marmoset, and macaque cerebellar ST and scRNA-seq data are accessible online at http://db.cngb.org/stomics/cbmsta.

## Code availability

The code of data processing and STACAME is accessible via this link: https://github.com/zhanglabtools/STACAME/.

## Acknowledgements

This work has been supported by the National Natural Science Foundation of China (no. 12501695 to B.Z., no. 12471350 to S.Q.Z., nos. 32341013, 12326614 to S.Z), the Hebei Natural Science Foundation (no. A2025203003 to B.Z.), the Science and Technology Commission of Shanghai Municipality (no. 23JC1401000 to S.Q.Z.), and the CAS Project for Young Scientists in Basic Research (no. YSBR-034 to S.Z.).

## Author contributions

S.Z. conceived the project. B.Z. and X.Z. developed and implemented the STACAME algorithm and the analysis. B.Z., X.Z., S.Q.Z. and S.Z. validated the methods and wrote the manuscript. S.Q.Z. and S.Z. supervised the project. All authors read and approved the final manuscript.

