## Supplemental Figures and Tables for "Deciphering shared and divergent tissue architectures from cross-species spatial transcriptomics"

### Supplementary Figures

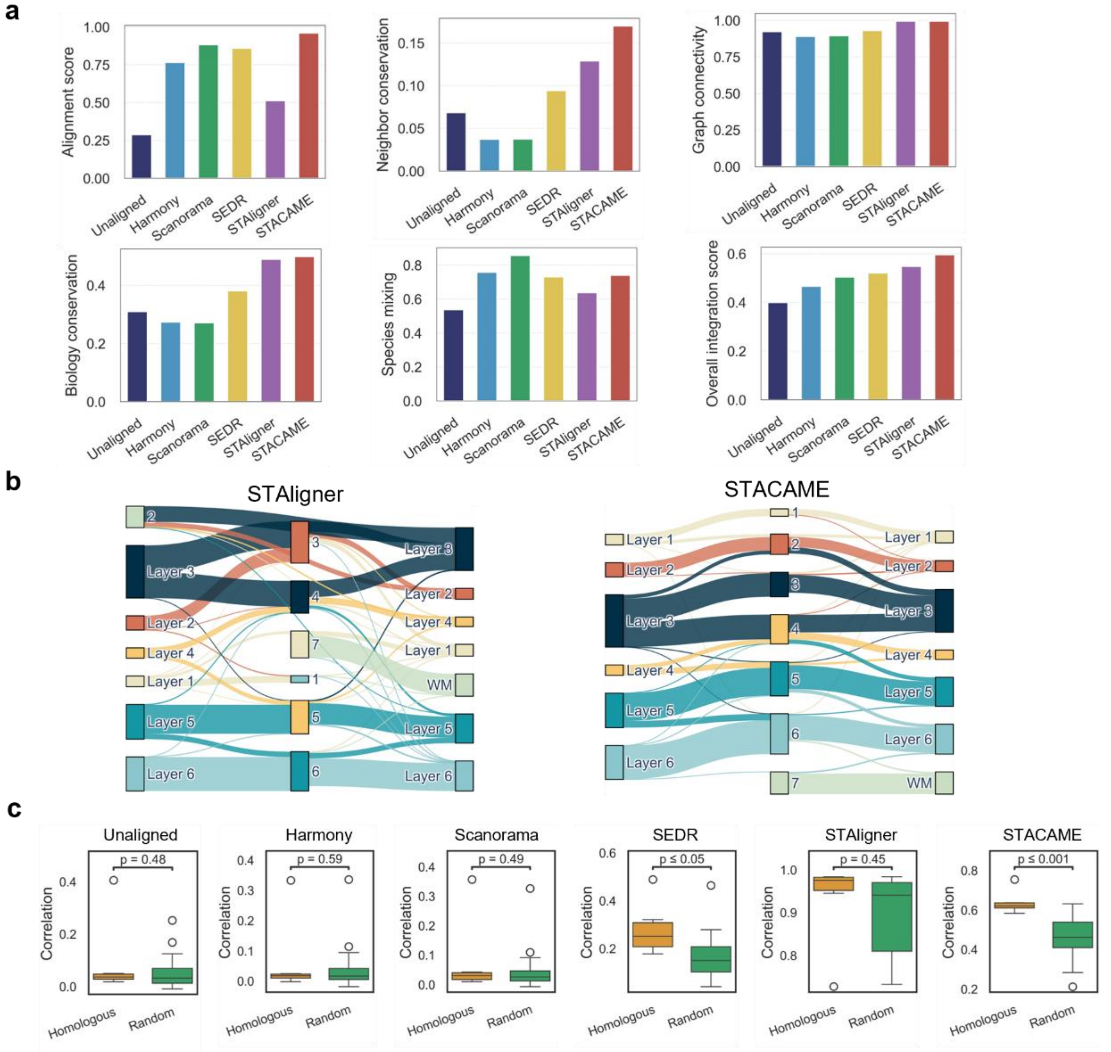

**Supplementary Figure 1. Additional performance analysis of STACAME on the mouse and macaque DLPFC datasets. a**, Comparison of embeddings' metric values computed with baseline methods and STACAME. **b**, Sankey plots of alignment among mouse regions, shared clusters and macaque regions for STAligner[1] and STACAME embeddings, respectively. The line width between two nodes is the spot proportion. **c**, The correlation of spots in homologous cortex layers between macaque and human, compared to random spot correlations. The center line, box limits, and whiskers denote the median, upper and lower quartiles, and 1.5x interquartile range, respectively, in the boxplot. The P-value for a two-sided hypothesis test whose null hypothesis is that the means of the two groups are equal, using Welch's t-test with Bonferroni correction for multiple comparisons.

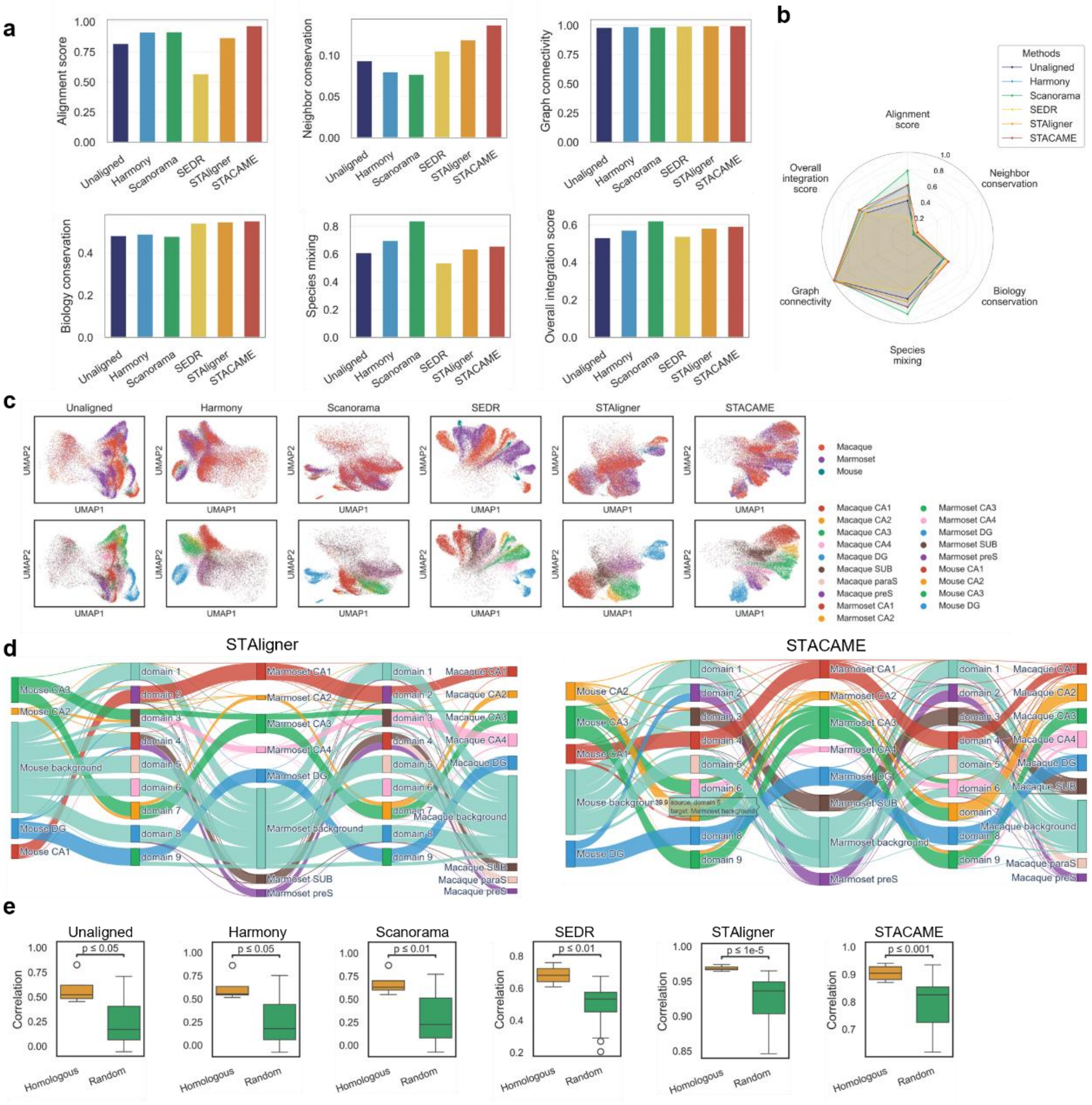

**Supplementary Figure 2. The integration performance of Harmony[2], Scanorama[3], SEDR[4], STAligner and STACAME on the mouse, marmoset and macaque hippocampus (Stereo-seq).** **a**, **b**, Comparison of embeddings' metric values. **c**, UAMP of mouse and macaque spot embeddings. **d**, Sankey plots of alignment among mouse regions, shared clusters and macaque regions for STAligner and STACAME embeddings. The line width between two nodes is the spot proportion. **e**, The correlation of spots in homologous regions between mouse and macaque, compared to random spot correlations. The center line, box limits, and whiskers denote the median, upper and lower quartiles, and 1.5× interquartile range, respectively, in the boxplot. The P-value for a two-sided hypothesis test whose null hypothesis is that the means of the two groups are equal, using Welch's t-test with Bonferroni correction for multiple comparisons.

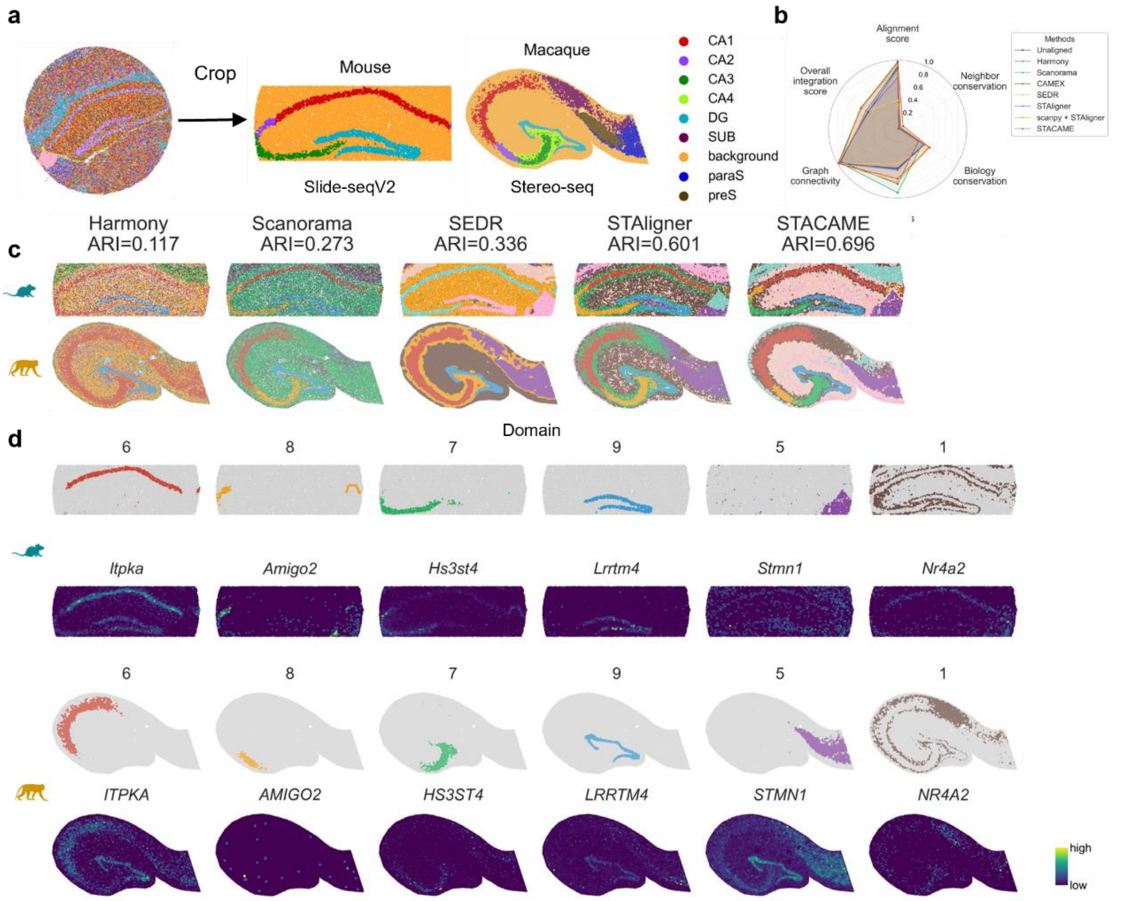

**Supplementary Figure 3. The integration performance of Harmony, Scanorama, SEDR, STAligner and STACAME on the mouse (Slide-seqV2) and macaque hippocampus (Stereo-seq). a**, Illustration of the mouse dataset sequenced by Slide-seqV2 and the macaque dataset sequenced by Stereo-seq is annotated by STAGATE5. **b**, Comparison of embeddings' metric values computed with baseline methods and STACAME. **c**, The spatial domain identified by STACAME and the other baseline methods. For the datasets of two species together, we run *mclust* (cluster number = 9) on the integrated embeddings and compute the ARI between manually annotated regions and clustered domains. **d**, Spatial mapping of identified hippocampus regions (CA1, CA2, CA3 and dentate gyrus) and marker genes. Mouse marker genes have been proven in previous references, and macaque marker genes here are the one-to-one homologous genes to the mouse.

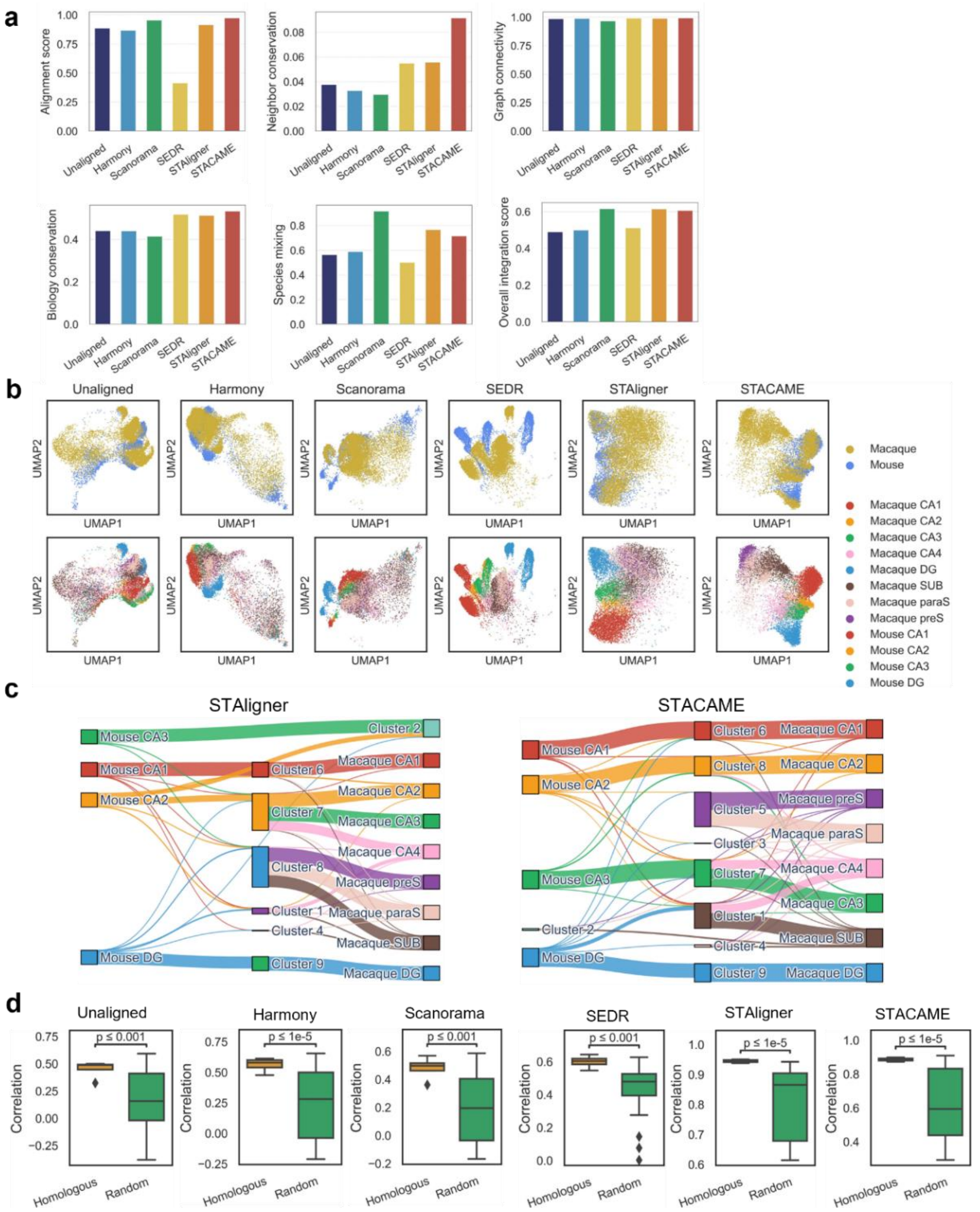

**Supplementary Figure 4. The integration performance of Harmony, Scanorama, SEDR, STAligner and STACAME on the mouse (Slide-seqV2) and macaque hippocampus (Stereo-seq).** **a**, Comparison of separate metrics. **b**, UAMP of mouse and macaque spot embeddings. **c**, Sankey plots of alignment among mouse regions, common clusters and macaque regions for STAligner and STACAME embeddings. The line width between two nodes is the spot proportion. **d**, The correlation of spots in homologous regions between mouse and macaque, compared to random spot correlations. The center line, box limits, and whiskers denote the median, upper and lower quartiles, and 1.5× interquartile range, respectively, in the boxplot. The *P*-value for a two-sided hypothesis test whose null hypothesis is that the means of the two groups are equal, using Welch's t-test with Bonferroni correction for multiple comparisons.

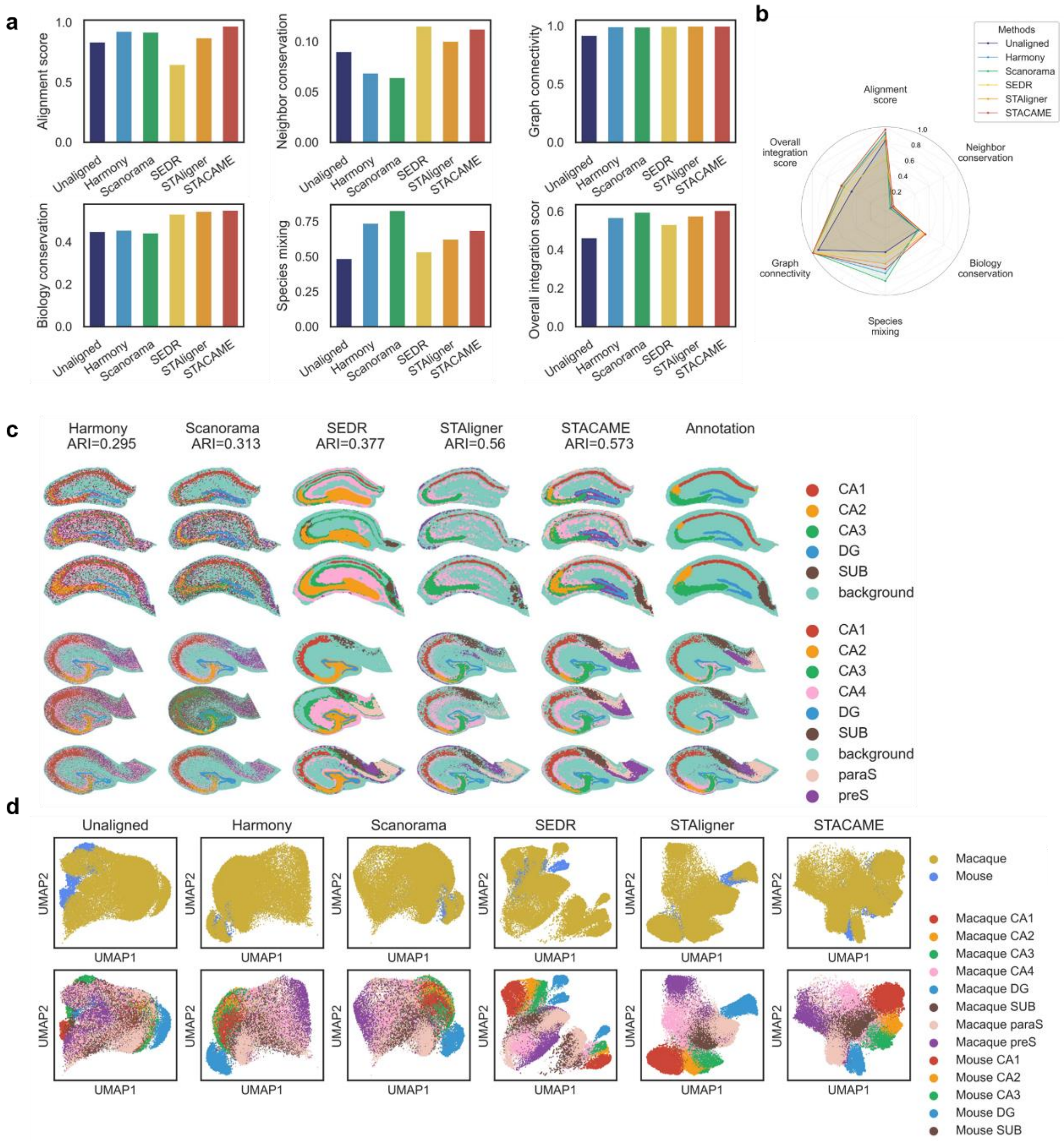

**Supplementary Figure 5. The integration performance of Harmony, Scanorama, SEDR, STAligner and STACAME on multiple spatial slices of mouse (Stereo-seq, T315, T319 and T323) and macaque hippocampus (Stereo-seq, T36, T38 and T42).** **a**, **b**, Comparison of separate metrics and overall integration score. **c**, Spatial domain identified by STACAME and the other baseline methods. For the two species' datasets together, we run *mclust* (cluster number = 16) on the integrated embeddings and compute the ARI between manually annotated regions and clustered domains. The spatial regions are annotated by STAGATE[2]. **d**, UAMP of mouse and macaque spot embeddings.

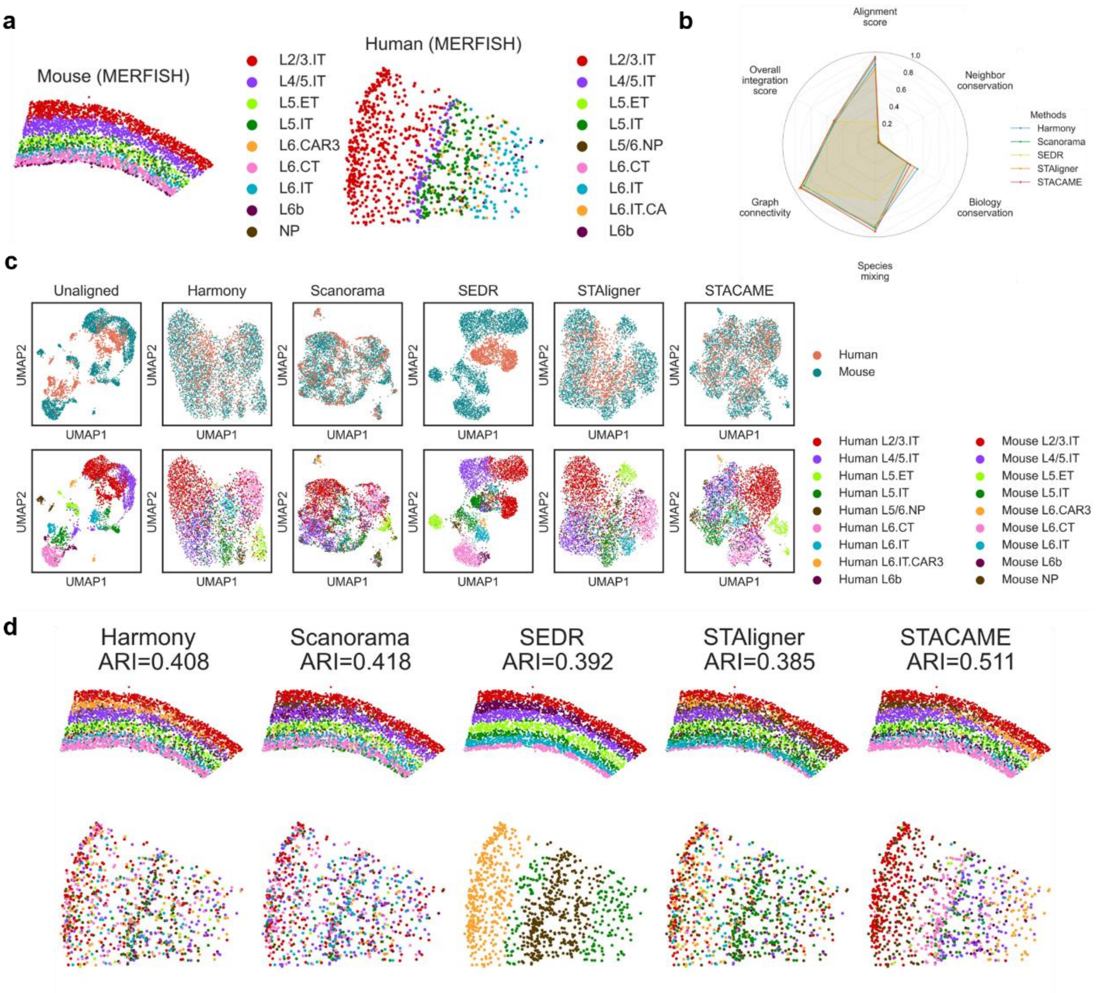

**Supplementary Figure 6. The integration performance of Harmony, Scanorama, SEDR, STAligner and STACAME on the mouse cortex (MERFISH) and human cortex (MERFISH).** **a**, Illustration of the mouse dataset and the human dataset sequenced by MERFISH, where cortex layers are annotated by their marker genes. **b**, Comparison of overall integration score. **c**, UAMP of mouse and macaque spot embeddings. **d**, The spatial domain identified by STACAME and the other baseline methods. For the two species' datasets together, we run mclust (cluster number = 9) on the integrated embeddings and compute the ARI between manually annotated regions and clustered domains.

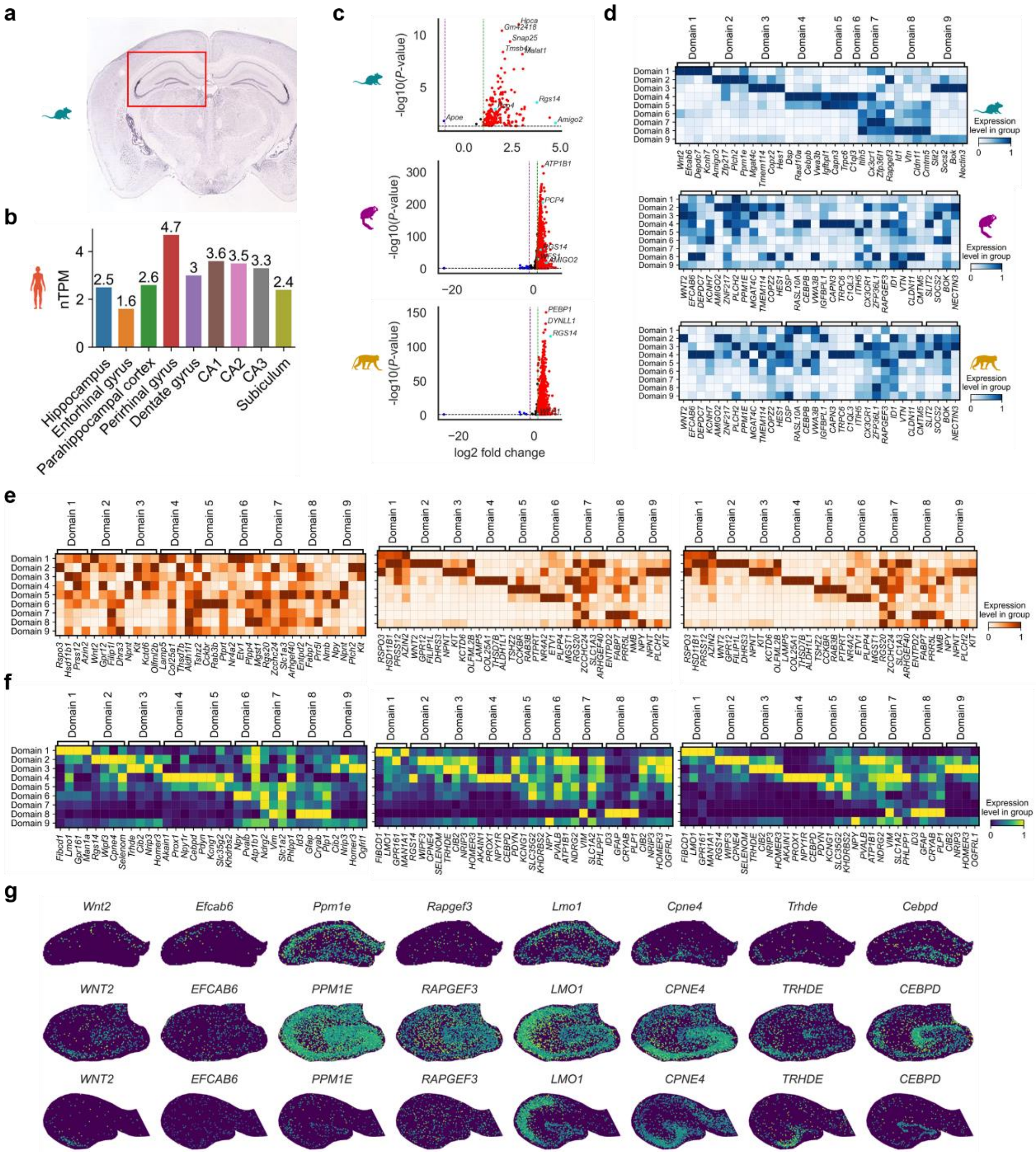

**Supplementary Figure 7. Additional analysis of cross-species consistent and divergent spatially variable genes (CCSVGs and CDSVGs). a**, Mouse *Amigo2* expression level on hippocampus in Allen mouse atlas ISH data[3]. **b**, Human *AMIGO2* protein expression level from the Human Protein Atlas[4]. **c**, Volcano scattering of mouse (left) and macaque (right) DEGs on CA2. The red dot denotes highly expressed genes (*P*-value < 0.05, fold change > 1); The blue dot represents lowly expressed genes (*P*-value < 0.05, fold change < -1); The black dot represents the other genes. We filtered out and marked known marker genes for CA2 if they are DEGs, and AMIGO2 is not significantly expressed in macaque CA2. We use a two-sided t-test to compare gene expression means between groups, with variance correction and Benjamini-Hochberg adjustment for multiple comparisons. **d**, Mean expression heatmap for shared spatial domains of mouse-specific DEGs whose homologous genes do not show significant differential expression for macaque. The genes are from the mouse DEGs, but not in the DEGs of marmoset or macaque (see **Methods**). **e**, Mean expression heatmap for common spatial domains of macaque-specific DEGs whose homologous genes do not show significant differential expression for mouse. The genes are from the macaque DEGs but not the mouse DEGs (see **Methods**). **f**, Mean expression heatmap of homologous DEGs showing high-level expression on common spatial domains for mouse, marmoset and macaque. **g**, Spatial expression patterns of representative CDSVGs and evolutionarily conserved spatially variable genes (CCSVGs) using original gene expressions.

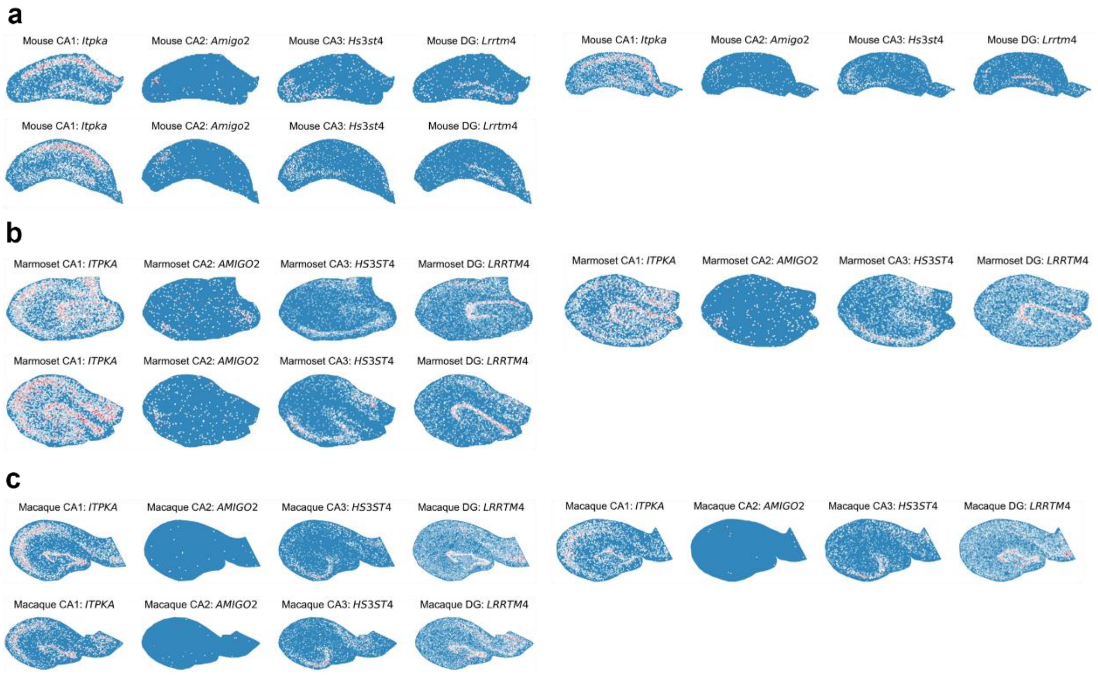

**Supplementary Figure 8. Expression of homologous marker genes (*Itpka*, *Amigo2*, *Hs3st4* and *Lrrtm4*) of the homologous brain regions in multiple spatial sections across the mouse, marmoset, and macaque hippocampus.** **a**, Mouse T315, marmoset T447 and macaque T36. **b**, Mouse T319, marmoset T458 and macaque T38. **c**, Mouse T323, marmoset T460 and macaque T42.

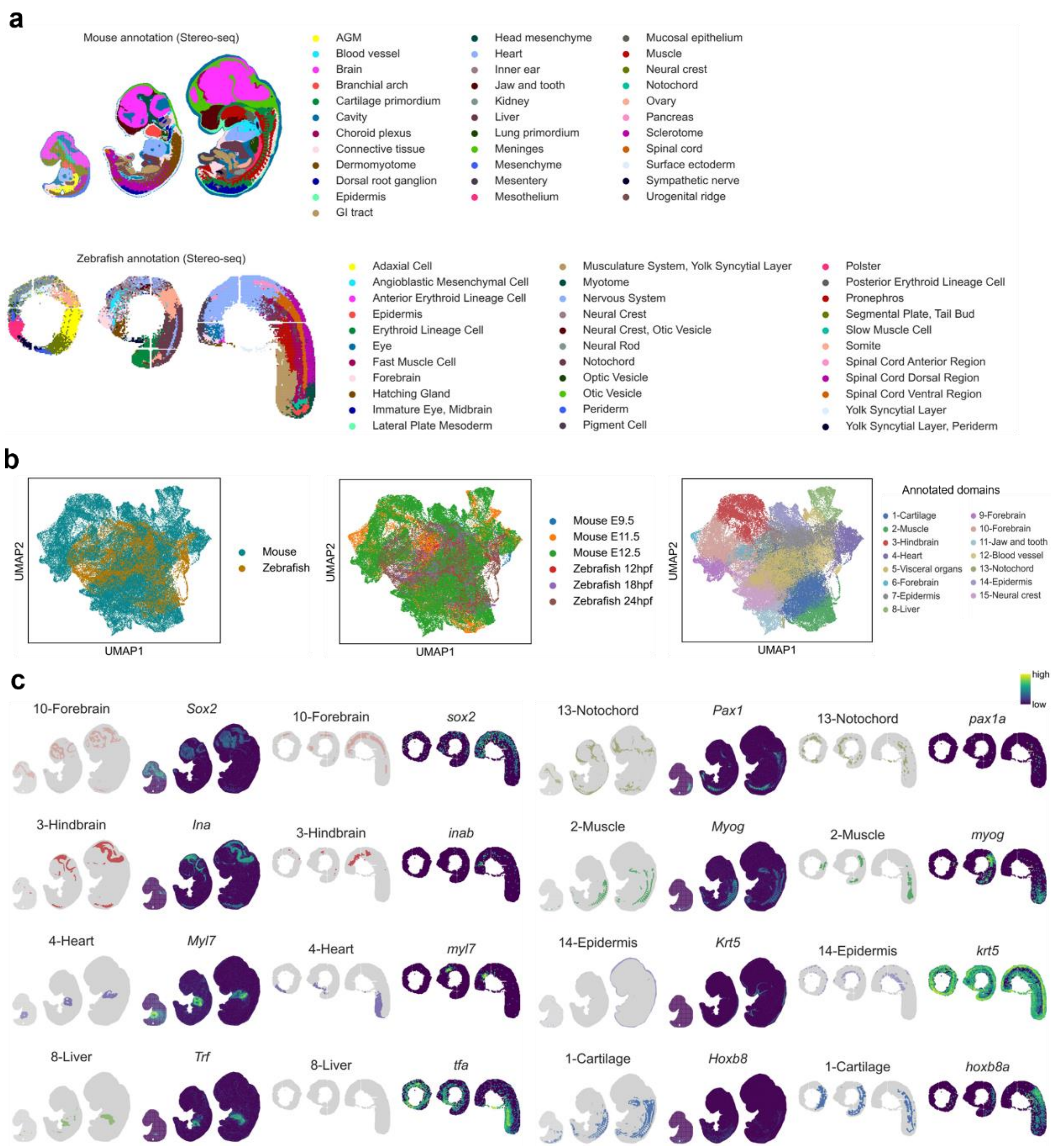

**Supplementary Figure 9. Integration of spatial transcriptomics across distantly related embryos (mouse and zebrafish) from multiple sections.** **a**, Manual annotation of mouse embryos at stages E9.5, E11.5, and E12.5 (Stereo-seq)[5] and zebrafish embryos at stages zf12, zf18, and zf24 (Stereo-seq)[6]. **b**, UMAP of spot embeddings labeled by species (left), slice names (middle) and identified domains (right). **c**, Spatial expression maps of 8 cross-species domains and their corresponding tissue marker genes identified by STACAME. Clustering was performed using *mclust* with 15 clusters.

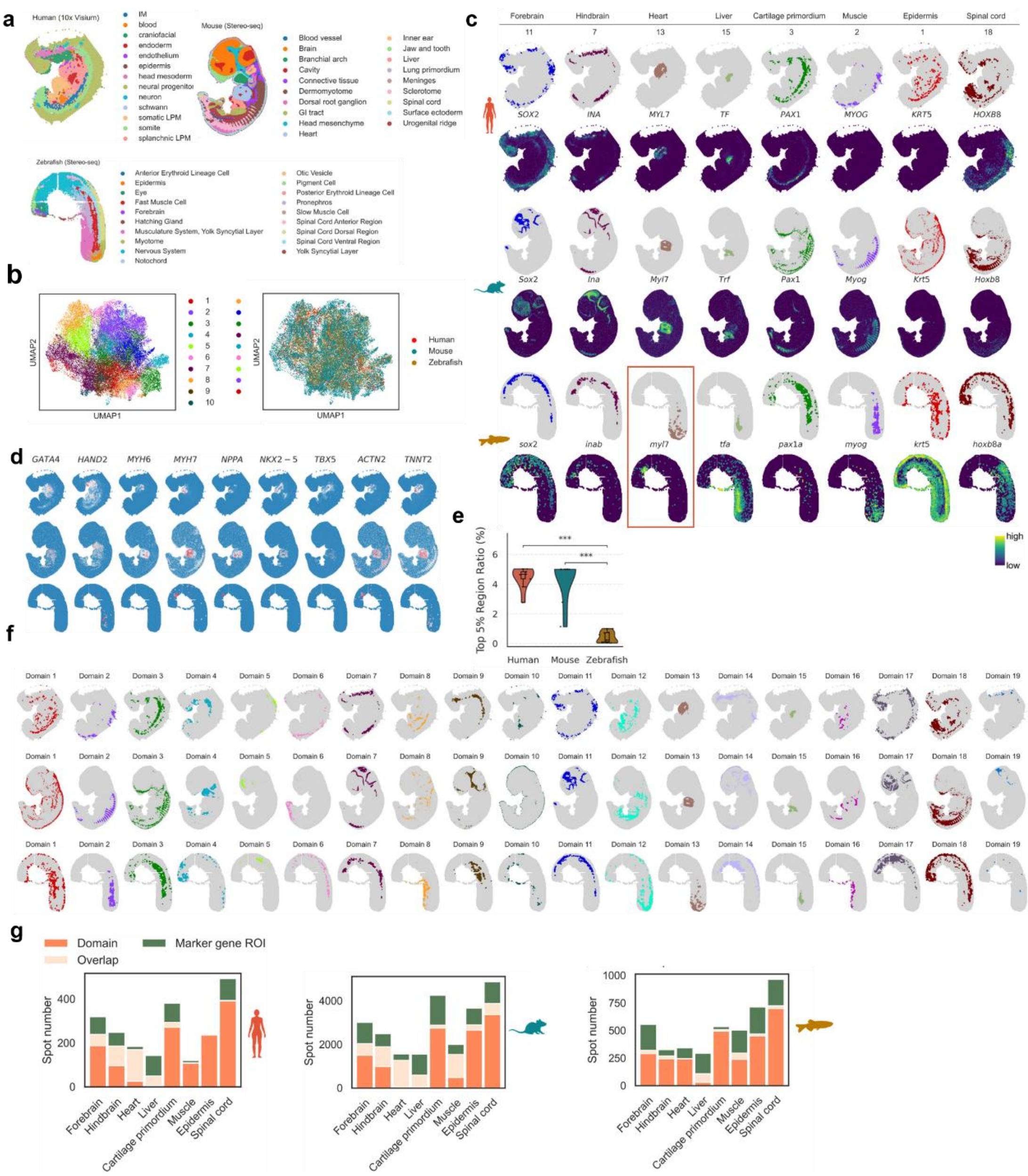

**Supplementary Figure 10. Integration of phylogenetically distant species' embryos. a**, Manually annotated human embryo at Carnegie stages (CS) 13 (10x Visium)[7], mouse embryo at stage E11.5 (Stereo-seq)[5] and zebrafish embryo at stage zf24 (Stereo-seq)[6]. **b**, UMAP of spot embeddings and identified shared domains across species. **c**, Cross-species domains identified via STACAME and expression spatial maps of corresponding tissues' marker genes. The clustering method is *mclust* with cluster number 19 (i.e., the annotated number of the mouse embryo). **d**, **e**, Analysis of the cause that STACAME failed to identify the zebrafish heart domain accurately.. **d**. The top 5% percentile high-expressing region ratio on the slice of 9 biomarkers, displays that the zebrafish embryo slice (24hpf) contains a significantly smaller heart domain (mouse vs zebrafish: *P*-value = $2.718 \times 10^{-4}$; human vs. zebrafish: *P*-value = $4.038 \times 10^{-4}$, with Mann-Whitney-Wilcoxon test two-sided). **e**, Spatial map of the 9 markers of heart genes, shared by human, mouse and zebrafish. Heart marker genes, such as *hand2*, display high expression levels in the STACAME-identified heart-corresponding domains. **f**, All the 19 identified homologous domains among three embryos via STACAME using multiple-to-multiple gene orthologs. **g**, The spot number in each common domain, marker gene highly expressed regions (ROI) and their overlap spot number for each species. The marker gene ROI consists of the top highly expressed spots (n equals the spot number in each common domain for the species), or the spots express higher than the 95th percentile if the minimum value of n top highly expressed spots equals zero.

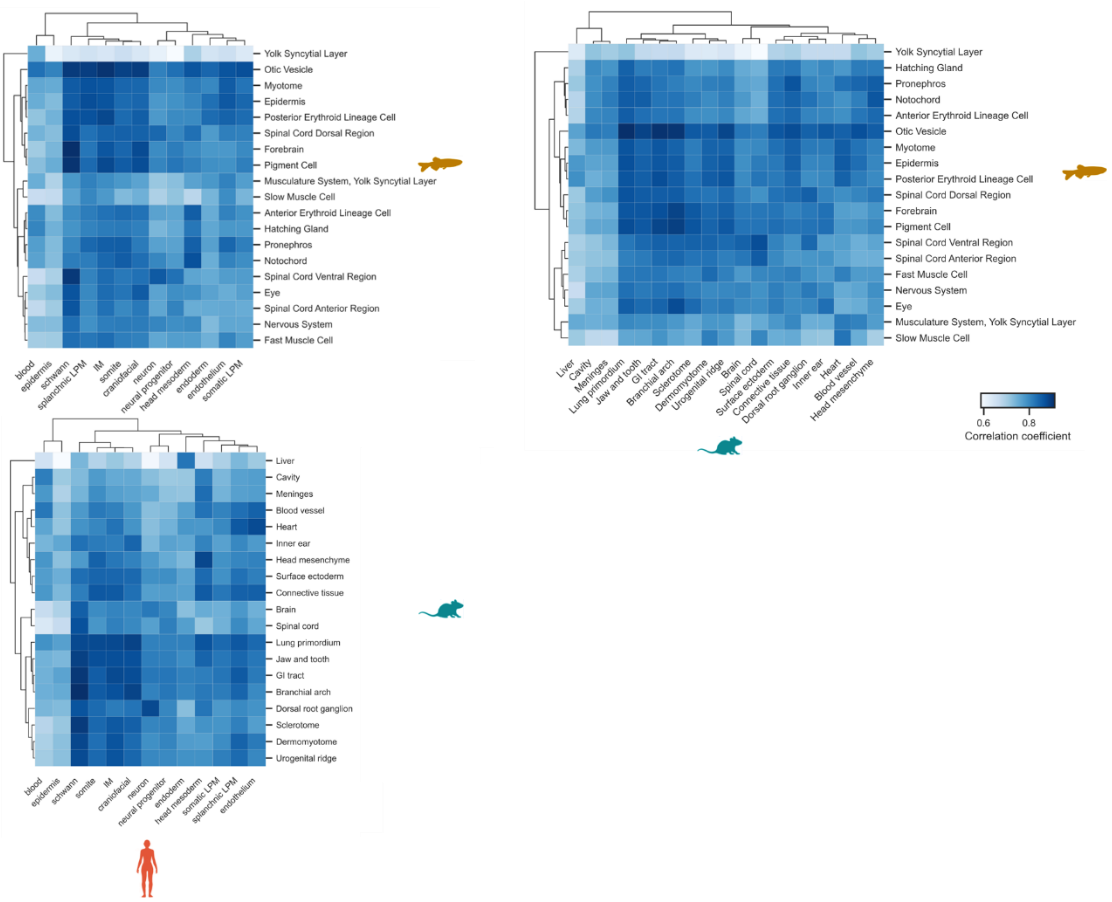

**Supplementary Figure 11. The heatmaps of the averaged Pearson correlation between any two annotated tissues among human, mouse and zebrafish embryos in Supplementary Figure 10.**

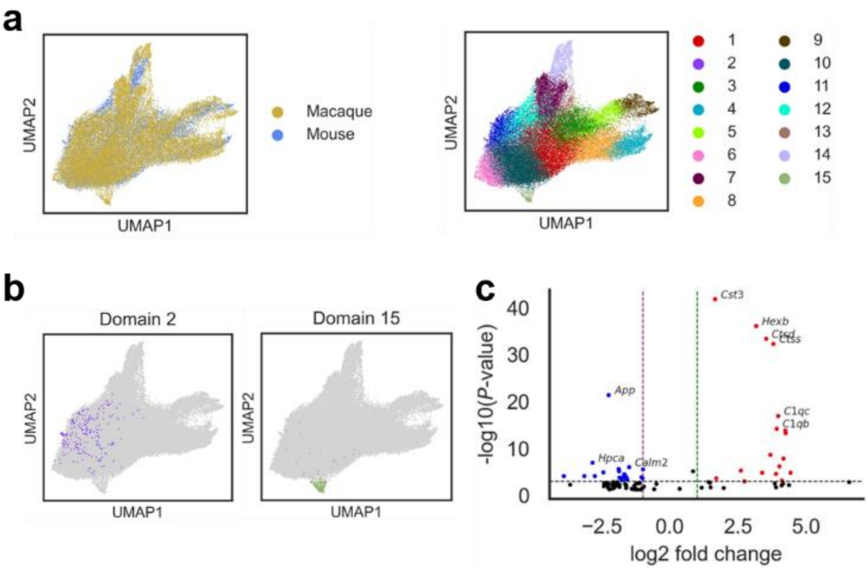

**Supplementary Figure 12. Additional results of integrating the normal macaque hippocampus ST slice and the mouse hippocampus ST slice in AD conditions.** **a**, UMAP of spot embeddings labeled by species and domains. **b**, UMAP presentation of mouse-specific domain 2 and macaque-specific domain 15. **c**, Volcano plot of DEGs between domain 2 of the mouse and the other domains.

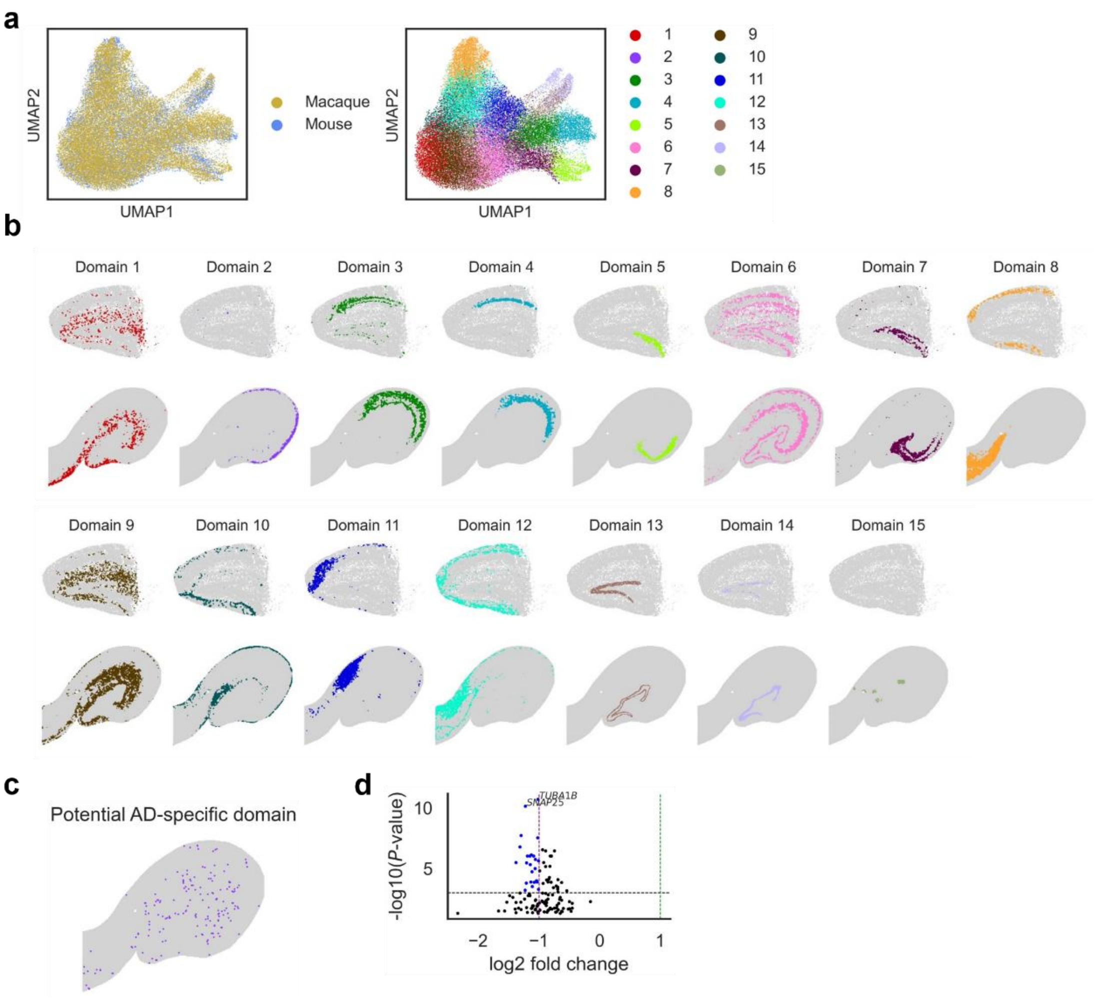

**Supplementary Figure 13. Illustration of the cause of Alzheimer's disease and determining the potential regions in the healthy macaque hippocampus using the integrated embedding of normal macaque hippocampus ST slice and mouse hippocampus ST slice of AD conditions (removing AD-related genes). a,** UMAP of embeddings produced by STACAME with domains assigned by *mclust*, where the integration experiment is conducted after removing AD-related genes in the mouse section. **b,** Spatial map of domains shared by mouse and macaque, where each domain corresponds to a pair of homologous regions. The AD domain has disappeared, demonstrating the differential expression of the removed AD-related DEGs. **c,** Macaque spots that correspond to the mouse AD domain in the embedding space without AD-specific genes as input. We run the MNN algorithm on embeddings to find them. d, Volcano plot of DEGs between macaque's AD-related domain and the other domains.

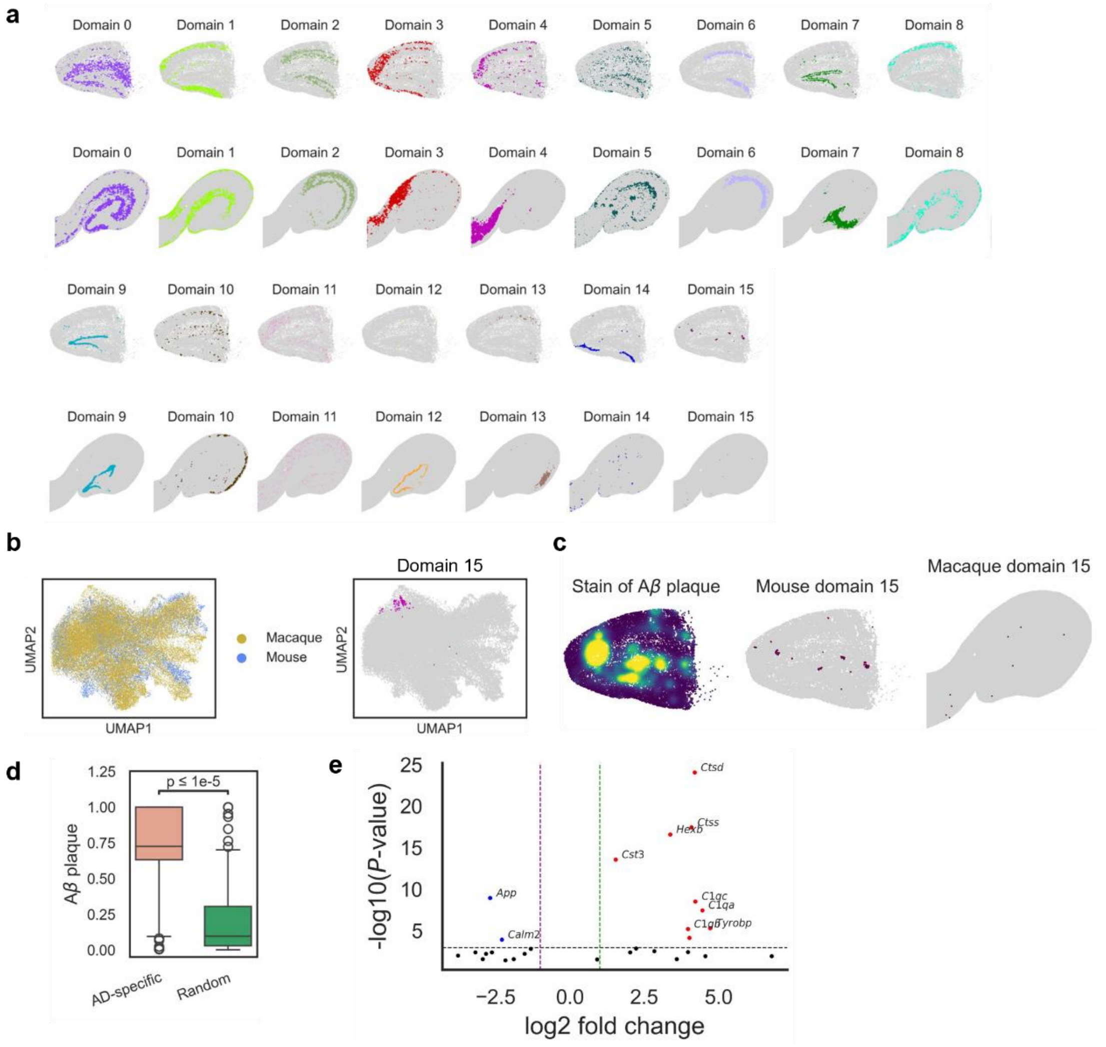

**Supplementary Figure 14. Additional results of Integrating normal macaque hippocampus ST slice and mouse hippocampus ST slice of AD conditions, clustering using *Louvain*.** **a**, Spatial domains assigned by *Louvain* on STACAME embeddings. **b**, UMAP presentation of mouse-specific domain 15. **c**, Adjacent tissue section showing Aβ plaque distribution (left), spatial map of domain 15 in mouse (middle), and in macaque (right). **d**. Comparison of amyloid-*β* (A*β*) plaque immunostaining intensity in mouse domain 2 versus randomly selected control spots (same number). Boxplots as in Figure 5; *P* value from Welch's *t*-test with Bonferroni correction. **e**, Volcano plot of DEGs between domain 15 of the mouse and the other domains.

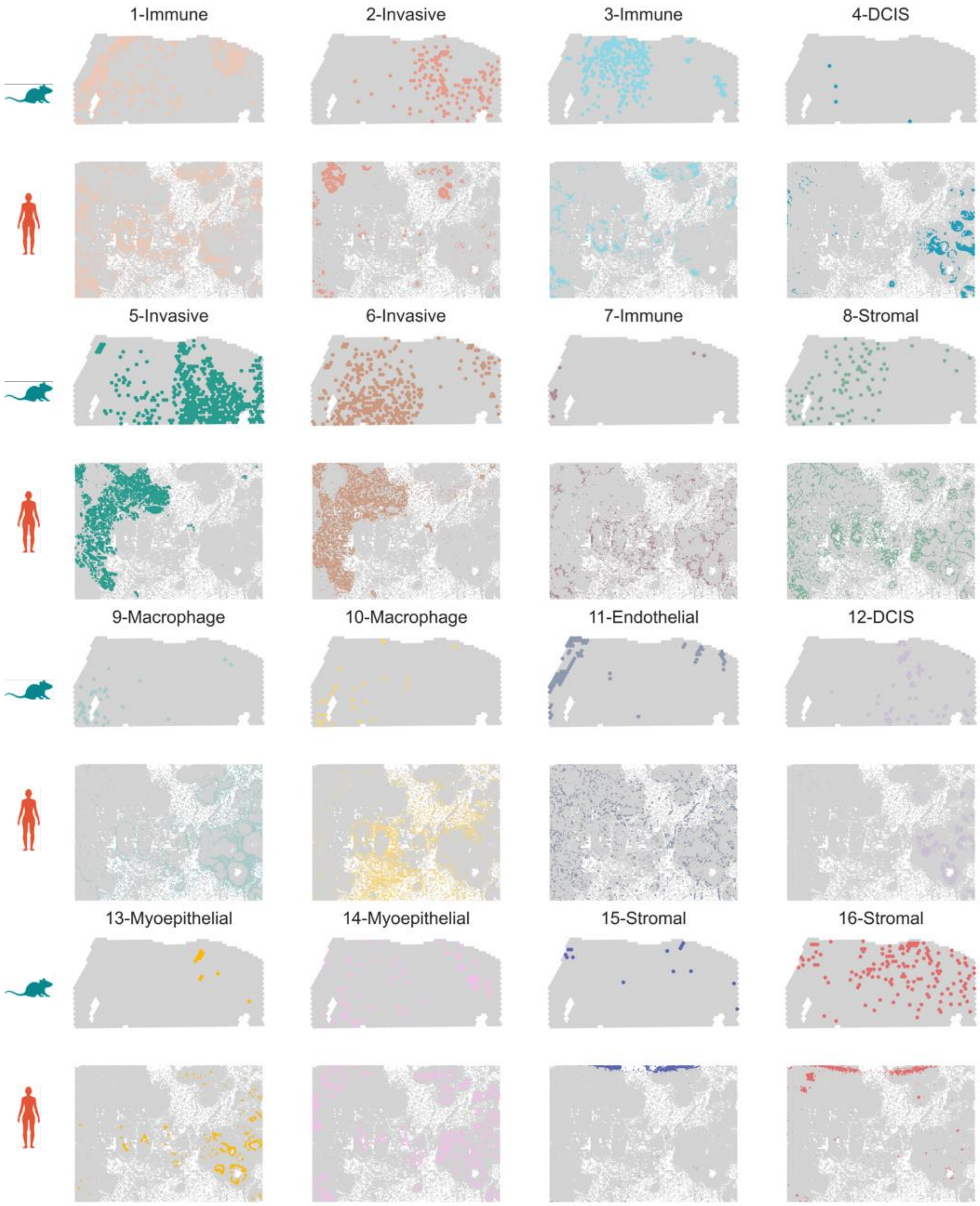

**Supplementary Figure 15. Cross-species domains identified by STACAME on the mouse and human breast cancer (BC) tissues profiled by 10x Visium and 10x Xenium, respectively.**

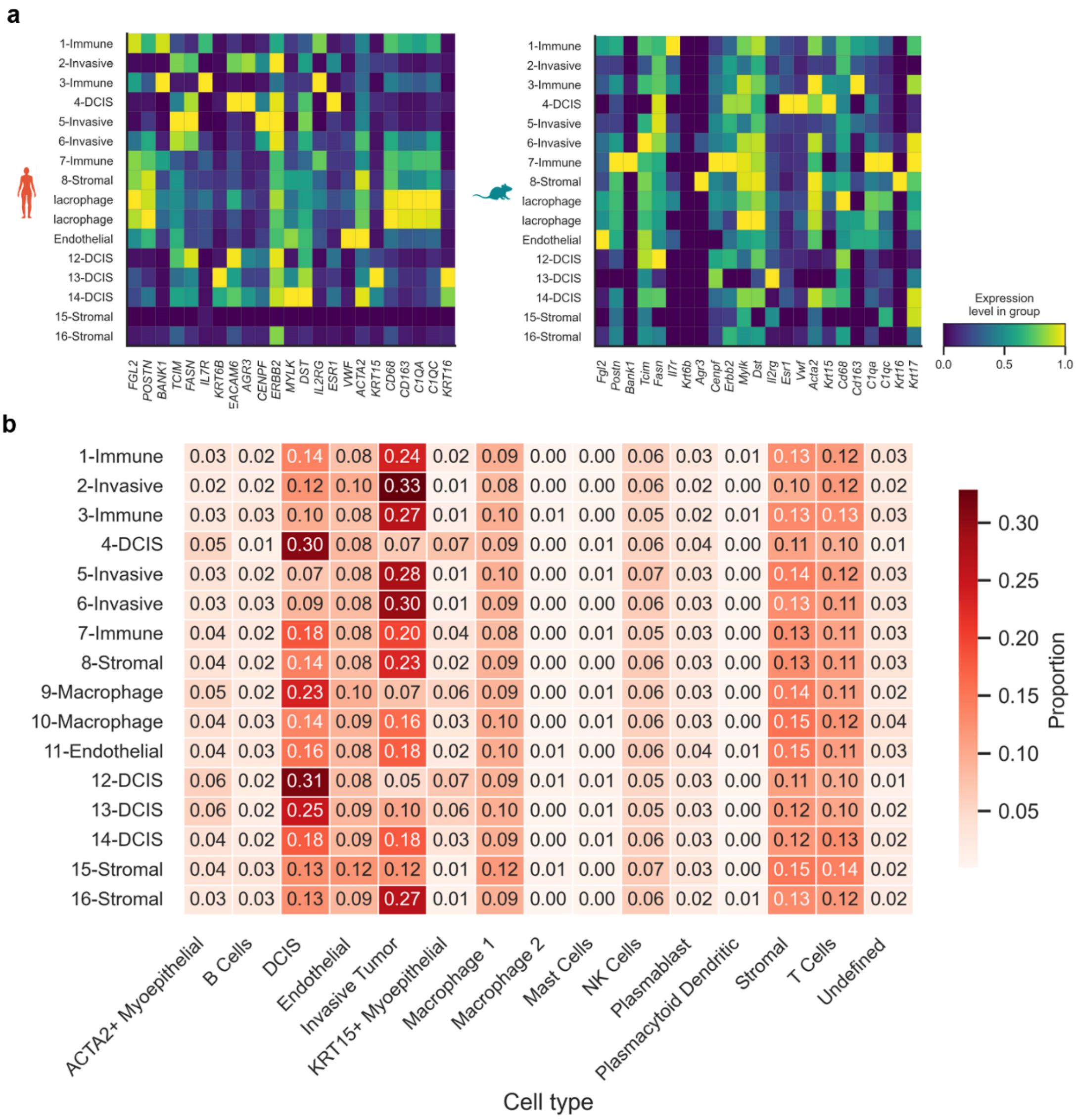

**Supplementary Figure 16. Additional results for cross-species breast cancer integration. a**, Mean expression of known marker genes on shared domains across mouse (right) and human (left). **b**, Cell type proportion of human domains.

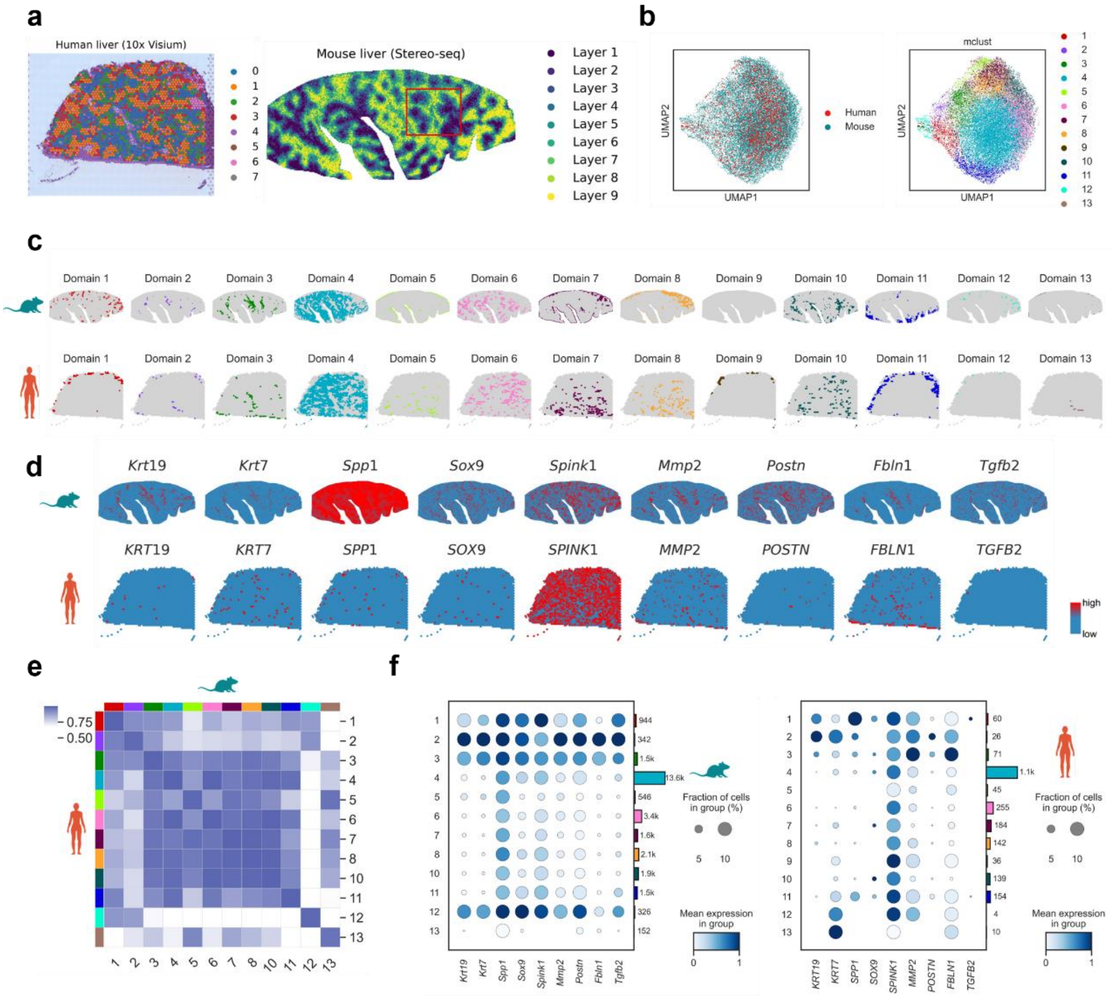

**Supplementary Figure 17. Integration of normal human liver ST section (Visium) and mouse liver ST section of cholestatic injury (Stereo-seq). a**, Human ST slice where labels are generated by clustering in the reference[8], and mouse ST slice where labels are manually annotated in the reference[9]. **b**, The UMAP of mouse and human liver spot embeddings produced by STACAME. **c**, Scattering of 13 Common domains identified by STACAME. **d**, Expression level of Important marker genes in the liver. The cholangiocyte domain (chol domain) was marked by *Krt19* and *Krt7* expression, whereas the LPLC domain was defined by co-expression of Spp1 and Sox9 coupled with the absence of *Krt19* and *Krt7*. **e**, The embedding average correlations among mouse and human liver domains. **f**, Dot plot of gene expression and spot percentages of important liver-related marker genes.

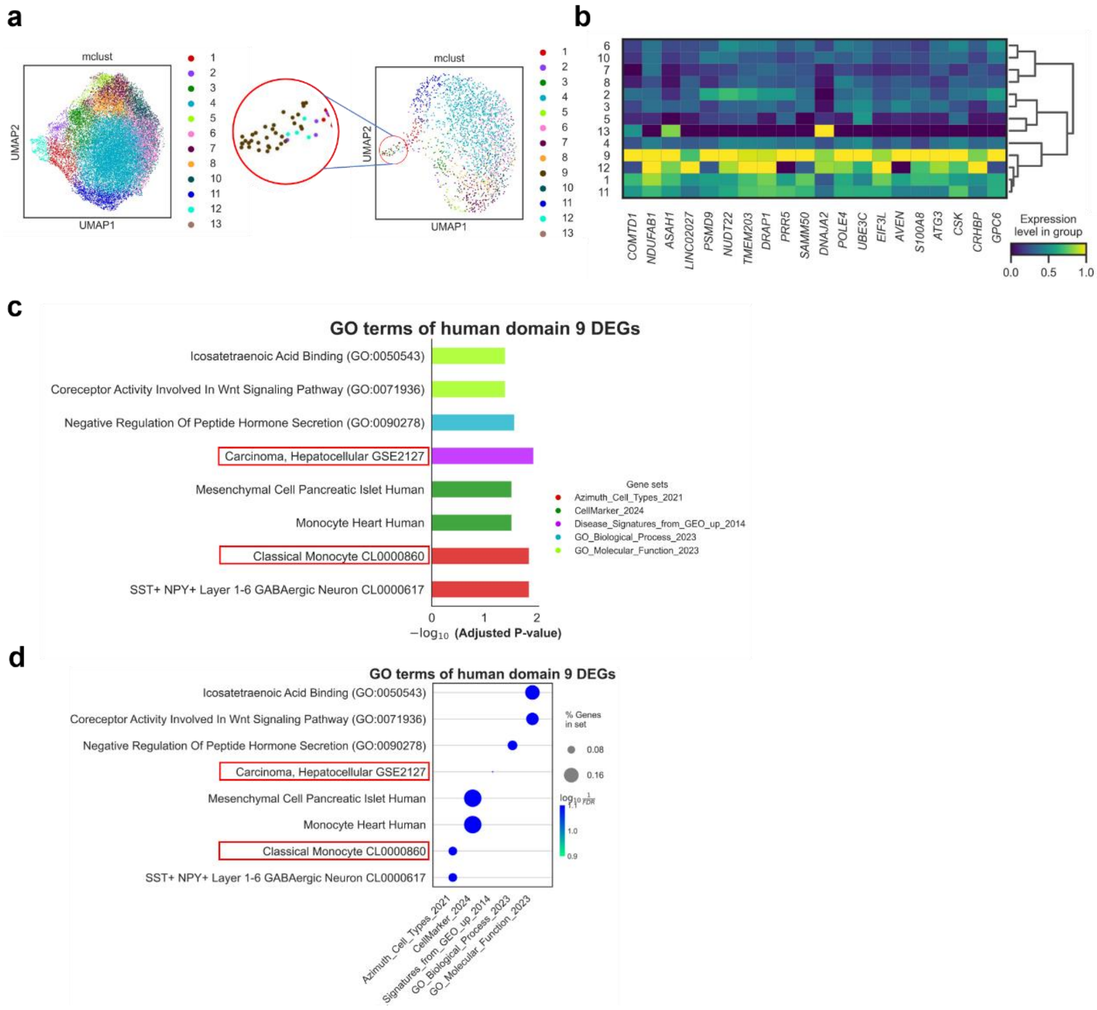

**Supplementary Figure 18. Analysis of the human liver-specific domain identified by STACAME. a**, UMAP of mouse and human STACAME embeddings, respectively. Domain 9 (red frame) only appears in human spots. **b**, The mean expression heatmap of the human DEGs of domain 9. The matrix data is normalized in the variable direction by subtracting the minimum and dividing each by its maximum for each gene. **c-d**, Mostly enriched GO terms of domain 9's DEGs within the human dataset measured by *P*-values. Hepatocellular Carcinoma and monocyte (red frame) probably indicate that this liver tissue is already in a mildly pathological state. In both bar plot (**c**) and dot plot (**d**), the *P*-value for a two-sided hypothesis test whose null hypothesis is that the frequency of genes in the input list belonging to a particular gene set is equal to the expected frequency by chance, using Fisher's exact test with Benjamini-Hochberg correction for multiple comparisons. Each row in the dot plot represents a different gene set that has been analyzed for enrichment. The size of each dot indicates the enrichment score for the corresponding gene set. The color of the dots indicates the negative logarithm of the *P*-value for a two-sided Fisher's exact test with Benjamini-Hochberg correction for multiple comparisons. The bar plot and dot plot are made by *gseapy* in Python.

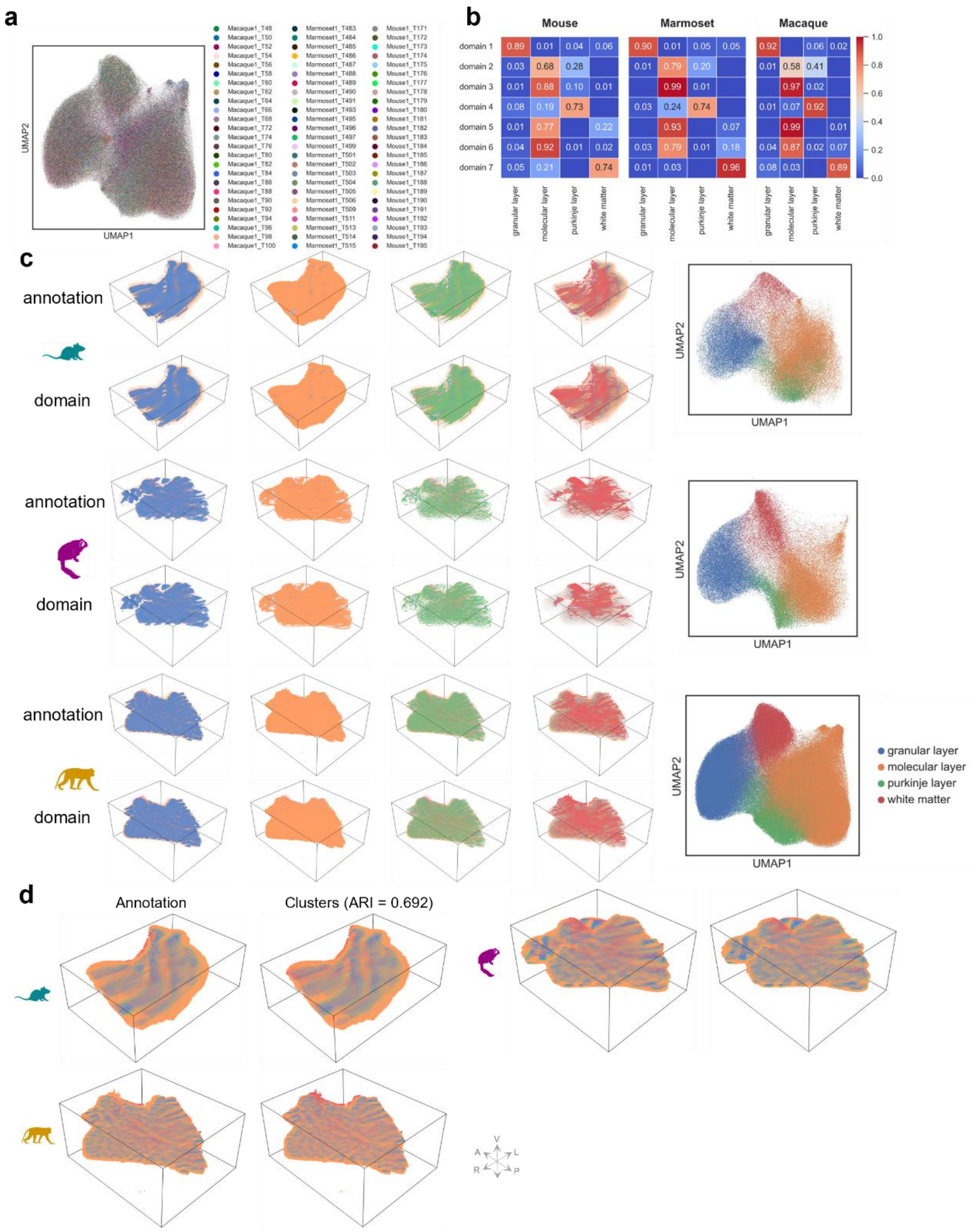

**Supplementary Figure 19. Additional analysis of cross-species integration of cerebellar from mouse, marmoset and macaque for more than ten million spots.** **a**, UMAP of STACAME spot embeddings labeled by slice names. **b**, Confusion matrices of domains and annotations for each species. **c**, 3D visualization of each annotated layer and its corresponding domain identified from STACAME embeddings for three species (left), and UMAP of spot embeddings labeled by annotated layers (right). **d**, The 3D visualization of the shared domains identified on STACAME embeddings and corresponding annotations. The 3D plot is generated with Spateo[10].

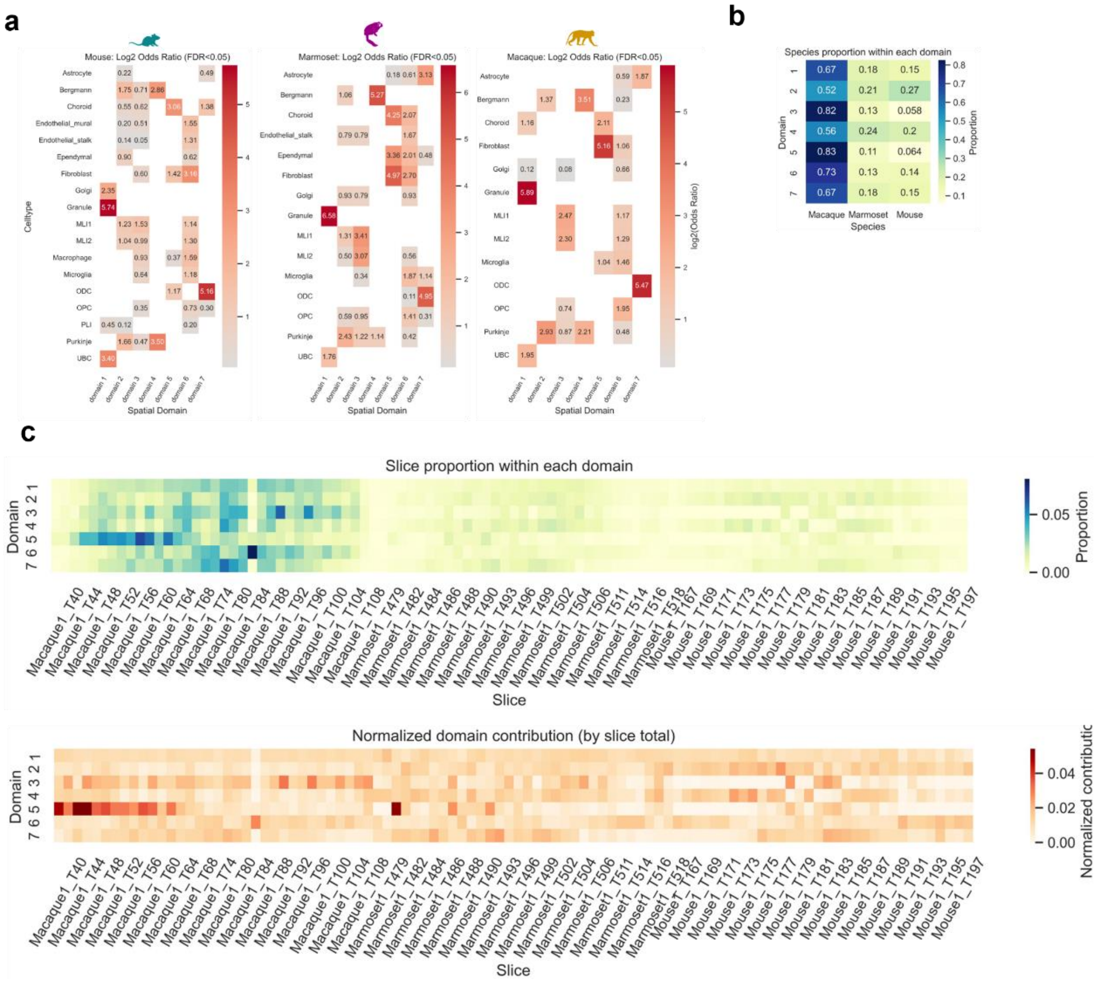

**Supplementary Figure 20. ST species and slices specificity on STACME domains and cell type specificity on domains. a**, Heatmaps show $\log_2$ odds ratios from one-sided Fisher's exact tests evaluating the association between individual cell types and spatial domains (domain 1–7) in each species. Only associations with false discovery rate (FDR) < 0.05 after Benjamini–Hochberg correction are displayed; non-significant entries are masked in white. Positive values (red) indicate significant enrichment of a cell type within a domain, while negative values (blue) reflect significant depletion. The divergent 'coolwarm' colormap is centered at zero to emphasize the directionality of association. Similar enrichment patterns across species suggest a conserved cellular architecture of spatial niches in the cerebellum. **b**, Heatmap of domain constitution of three species reveals three species-specific domains (2, 3 and 5). **c**, Heatmap of unnormalized (top) and normalized (bottom) domain constitution of each slice in three species reveals domain-specific slices.

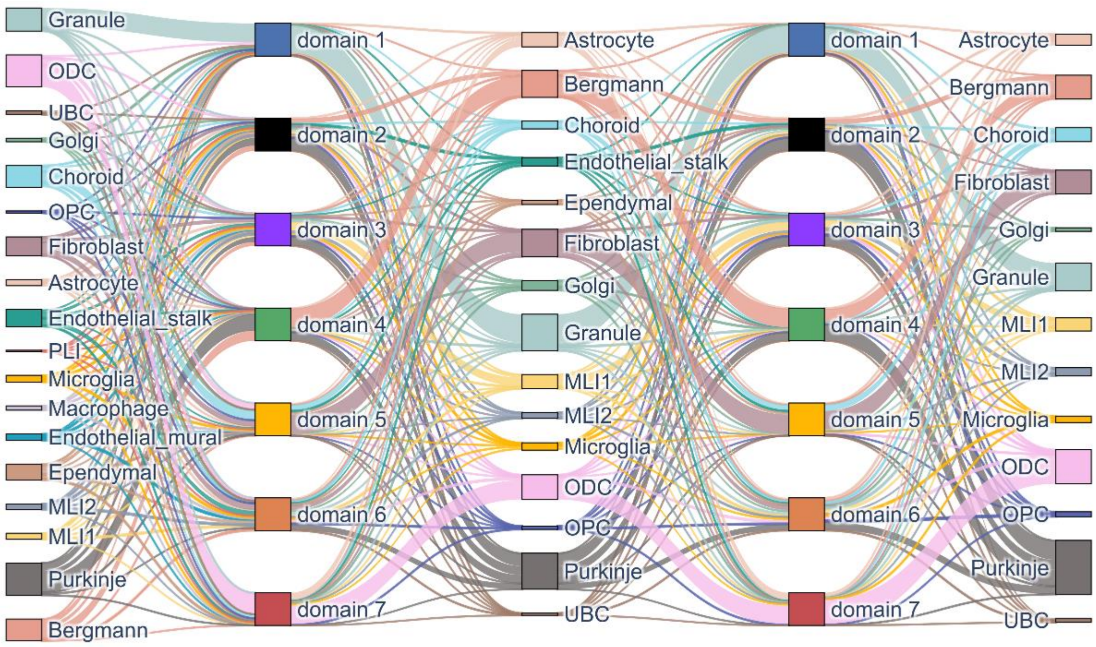

**Supplementary Figure 21. Sankey diagram depicting the flow of cell types through spatial domains across three species.** Each vertical column represents a layer in the cross-species mapping: mouse cell types → shared spatial domains (domain 1) → marmoset cell types → conserved spatial domains (domain 2) → macaque cell types. The width of each link is proportional to the relative abundance (normalized per domain) of a given cell type within its associated domain. Colors correspond to major cerebellar cell classes. This visualization highlights the preservation and divergence of cellular composition across evolutionarily conserved spatial niches in the cerebellum.

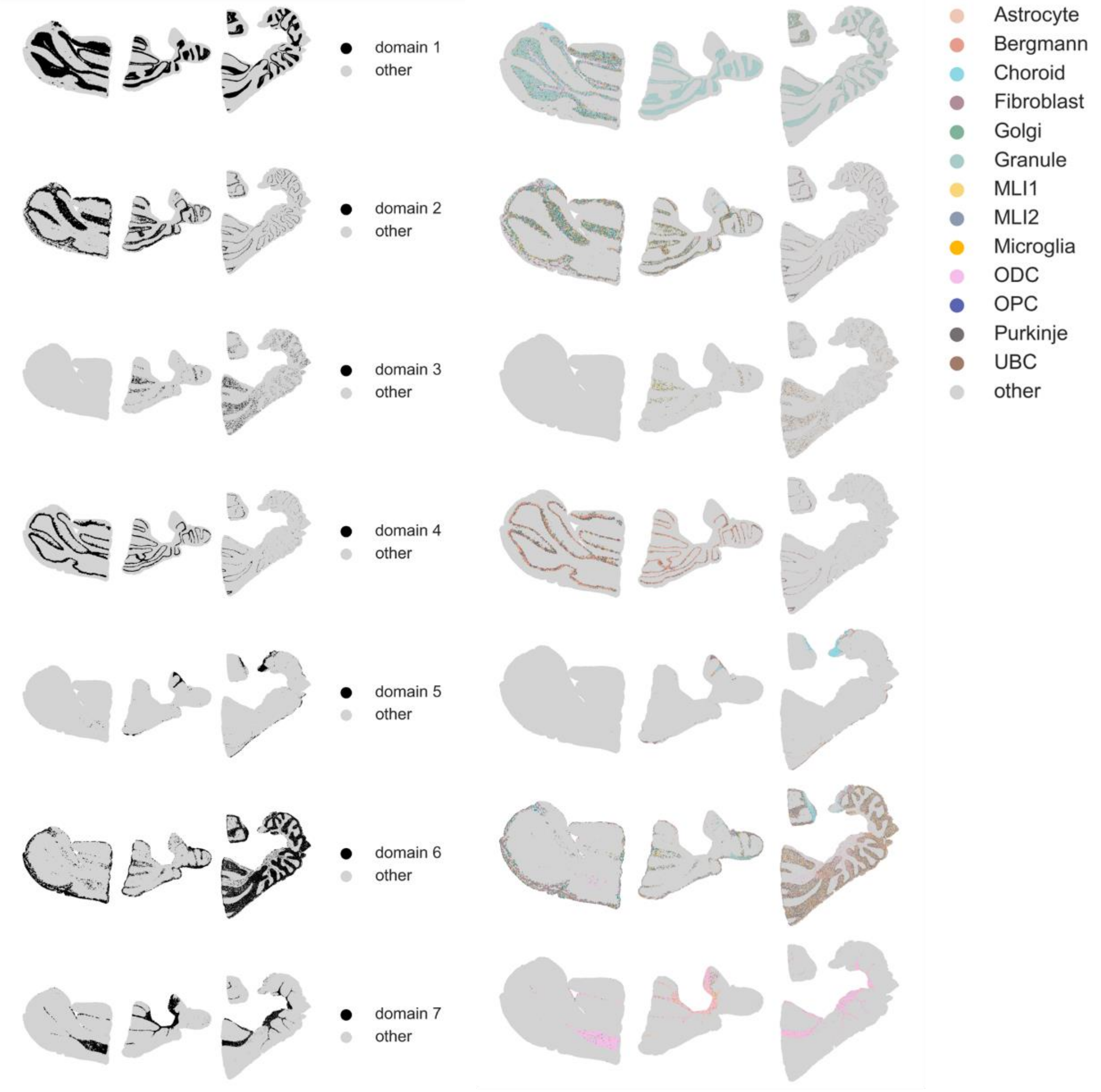

**Supplementary Figure 22. The spatial map of the three species-specific domains (T175, T490 and T62 for mice, marmosets and macaques, respectively) and their cell-type compositions.**

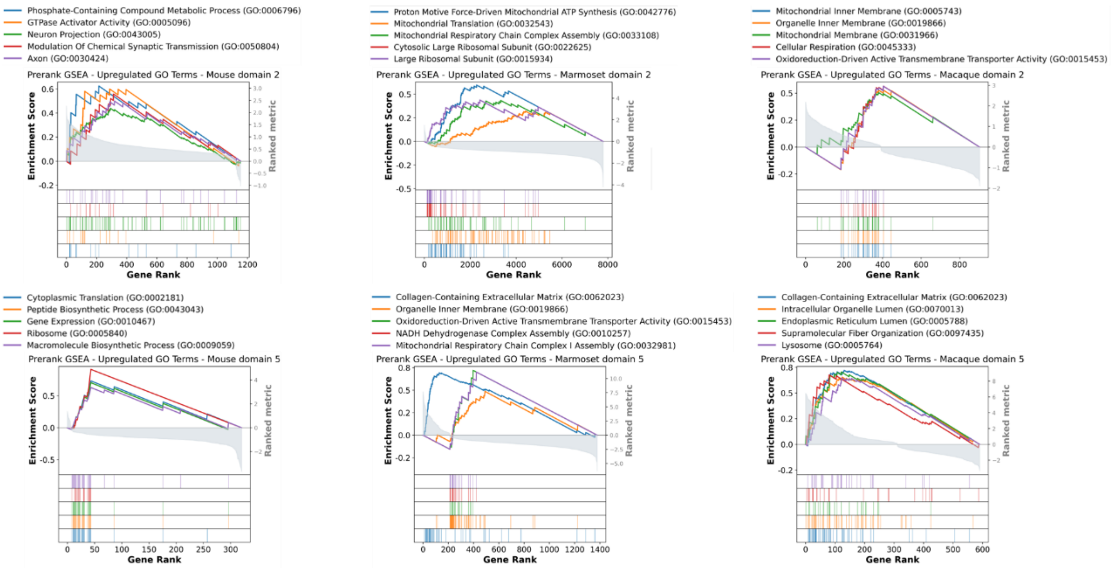

**Supplementary Figure 23. The gene rank plots of upregulated GO terms of domains 2 and 5.** The top GO terms display high similarities among species.

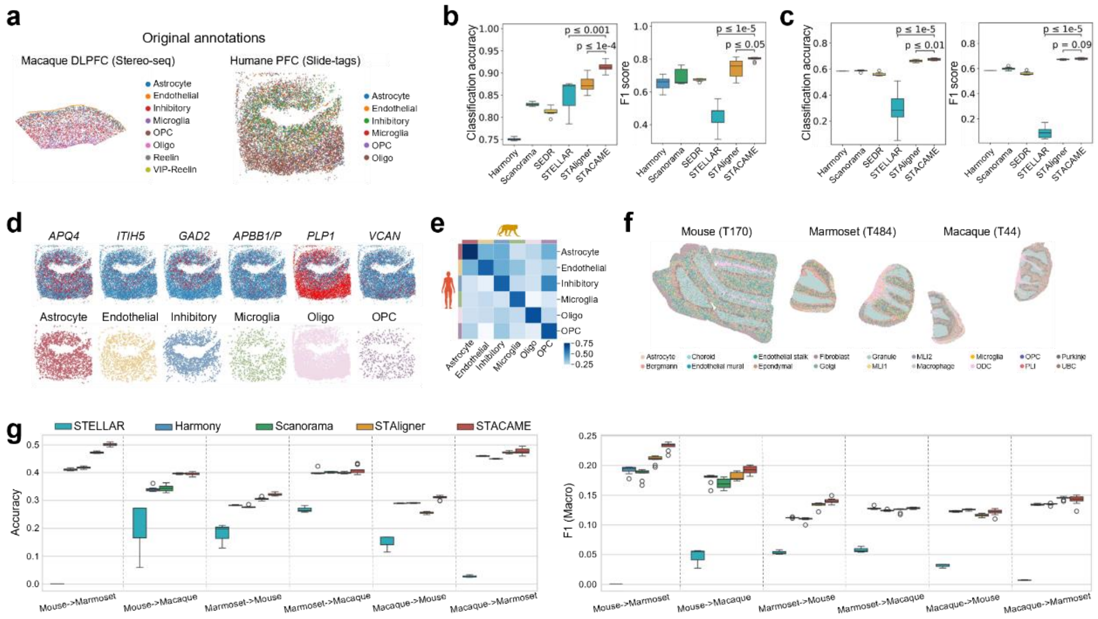

**Supplementary Figure 24. Cross-species prediction of neural cell types using STACAME embeddings.** **a**, Manually annotated cell types in macaque DLPFC and human prefrontal cortex (PFC, Slide-tags)[11]. **b, c**, Accuracy and macro F1-scores for predicting human PFC cell types using models trained on macaque DLPFC embeddings (**b**), and vice versa (**c**), using a two-layer fully connected neural network. Experiments were repeated five times with different random seeds. Boxplots show median (center line), interquartile range (box), and 1.5× IQR (whiskers). *P* values (two-sided Welch's *t*-test with Bonferroni correction) compare STACAME against baselines. **d**, Spatial expression of known human cortical cell-type markers (top) versus corresponding STACAME-predicted cell types (bottom). **e**, Heatmap of average embedding correlations between homologous cell types in macaque and human PFC. **f**, Manually annotated cerebellar cell types in mouse, marmoset, and macaque. **g**, Cross-species cell-type prediction performance (accuracy and macro F1-score) using embeddings from the other two species. STELLAR results are averaged over three runs.

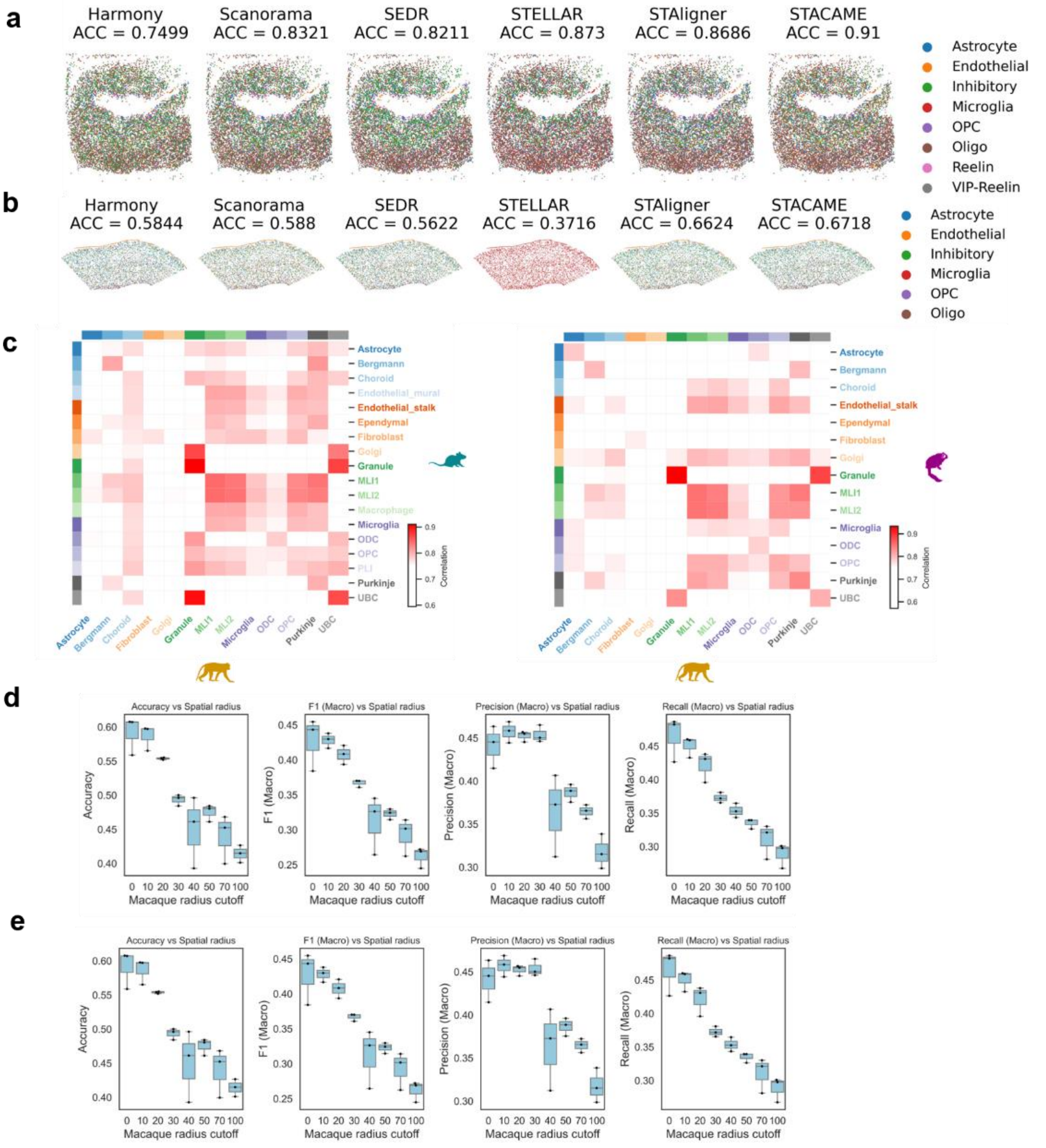

**Supplementary Figure 25. Additional results of cross-species spatial cell-type prediction.** **a**, **b**, Spatial mapping of predicted cell types and accuracy between background labels and predicting cell type labels for each method, using macaque DLPFC slice as test dataset (**a**) and human PFC slice as test dataset (**b**). **c**, The heatmap of spot embedding averaged correlations among cell types, across annotated mouse, marmoset and macaque cerebellar slices (T170 for mouse, T484 for marmoset, and T44 for macaque). **d**, **e**, The performance of STACAME using different amounts of spot spatial neighbor information, with macaque DLPFC slice as test dataset (**d**) and human PFC slice as test dataset (**e**). The accuracy, F1 (macro), precision (macro) and recall (macro) decline as spatial information increases.

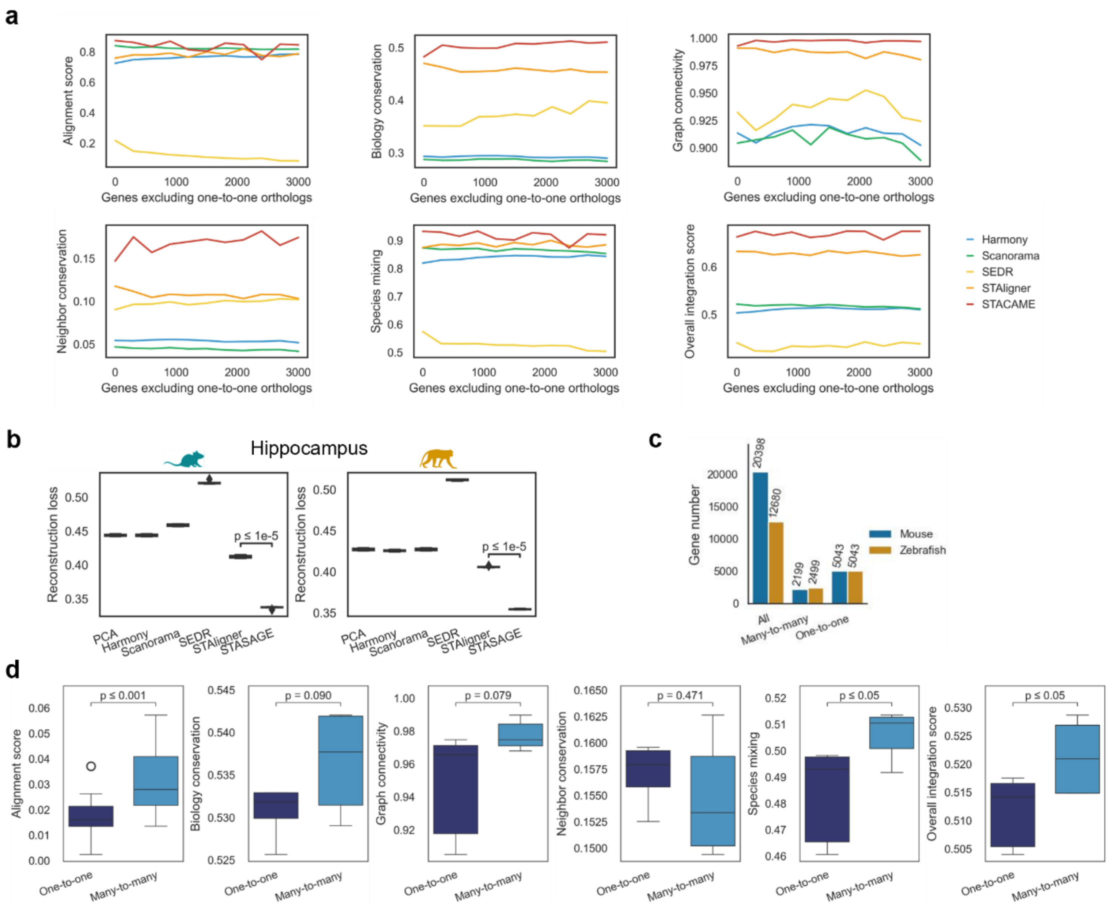

**Supplementary Figure 26. Ablation experiments for STACAME mechanisms. a**. Value change of integration evaluating metrics as non-homologous gene used in STACAME increases, with fixed hyper-parameters. **b**. The gene expression reconstruction loss of a trained two-layer FCN using the mouse and macaque embeddings (training: test = 8:2) for different methods. The training and test are repeated multiple times (n = 10) to reduce the effect of randomness. The center line, box limits, and whiskers denote the median, upper and lower quartiles, and 1.5x interquartile range, respectively. The P-value for a two-sided hypothesis test whose null hypothesis is that the means of the two groups are equal, using Welch's t-test with Bonferroni correction. **c**. The gene numbers, multi-to-multi homologous gene (exclude one-to-one) numbers, and one-to-one homologous gene numbers in the raw mouse (E11.5) and zebrafish (zf24) embryo datasets. **d**. Separate metrics and overall integration score value of embeddings using only one-to-one genes for building cross-species links and using both multi-to-multi and one-to-one genes. The two settings share the same hyper-parameters. We repeat 20 times with different random seeds (*P*-value = $8.031 \times 10^{-5}$). The center line, box limits, and whiskers denote the median, upper and lower quartiles, and 1.5x interquartile range, respectively. The *P*-value for a two-sided hypothesis test whose null hypothesis is that the means of the two groups are equal, using Welch's t-test with Bonferroni correction.

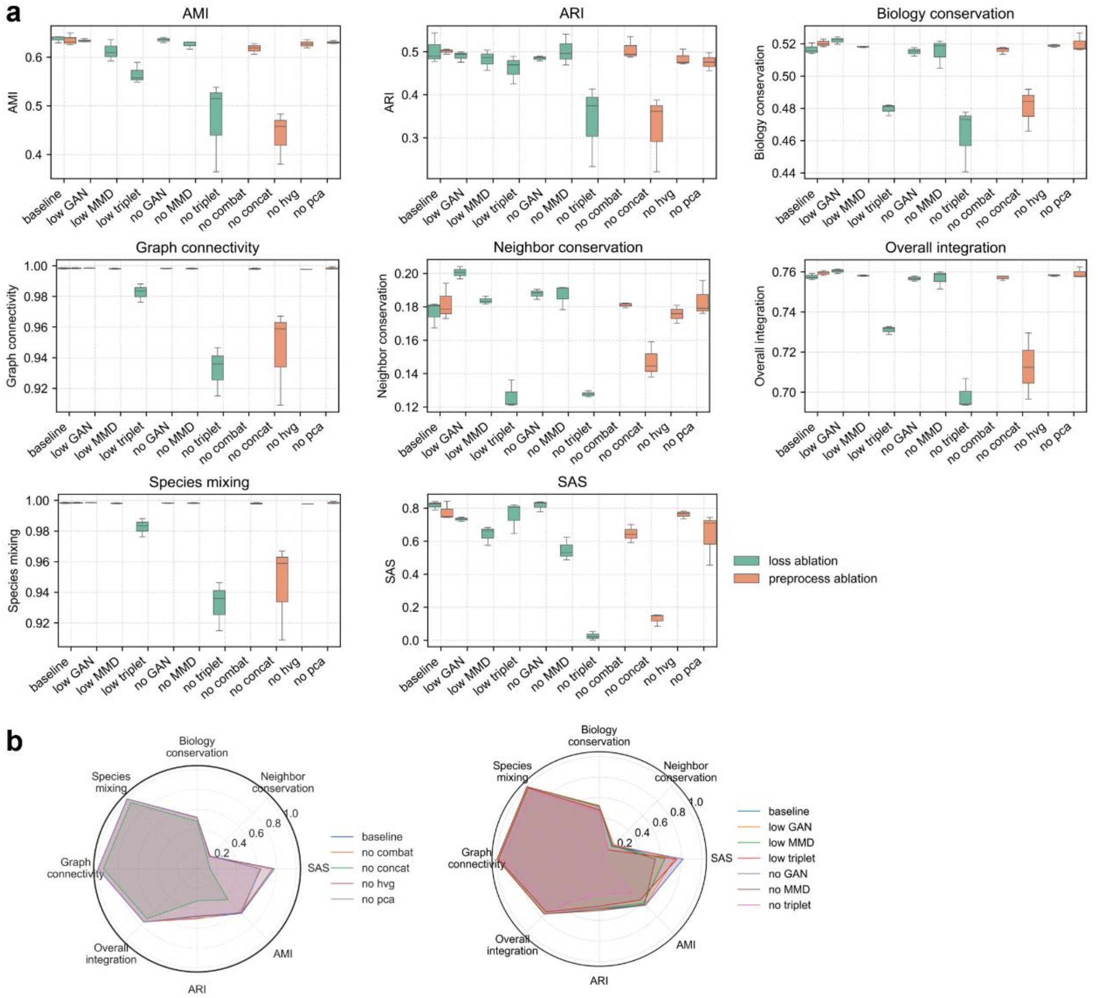

**Supplementary Figure 27. Ablation experiments for STACAME settings and hyper-parameters. a**, Boxplots of key integration and biological fidelity metrics across three independent runs for each ablation condition. Experiments are grouped by type: *loss_ablation* (top) varies the weighting coefficients of the maximum mean discrepancy (MMD), adversarial (GAN), and triplet loss terms in the total objective; *preprocess_ablation* (bottom) evaluates the impact of individual preprocessing steps—highly variable gene selection (HVG), combat-based batch correction, PCA projection before mutual nearest neighbors (MNN) alignment, and concatenated input construction. Metrics include Seurat alignment score (SAS), neighbor conservation, biology conservation (composite of spatial neighborhood preservation, mean average precision, and silhouette width), species mixing (average of graph connectivity and cross-species annotation alignment), overall integration, and clustering concordance (ARI/AMI). **b**, Radar plots summarizing the mean performance (averaged over three runs) of individual preprocessing steps (left) and each loss-ablation configuration (right). The baseline model employs balanced weights (MSD β=1, GAN β=10, triplet β=10); removing or down-weighting any component degrades specific aspects of integration or biological structure preservation, highlighting the complementary roles of distribution alignment (MSD), domain confusion (GAN), and intra-species discriminability (triplet loss) in achieving robust cross-species joint embedding.

### Supplementary Table

**Supplementary Table 1.** Summary of the spatial multi-species data used in this study.

| Exp id & Tissue | Species | Platform | Slice id | # of spots or cells | Related figures |
|---|---|---|---|---|---|
| 1-DLPFC | Human | 10x Visium | 151673 | 3,611 | Fig. 2 Fig. S1 Fig. S26a Fig. S27 |
| | Macaque | Stereo-seq | Part of T127 | 18,375 | |
| 2-Hippocampus | Mouse | Stereo-seq | T315 | 4,540 | Fig. 3 Fig. S2 Fig. S7 Fig. 26b |
| | Marmoset | Stereo-seq | T447 | 28,566 | |
| | Macaque | Stereo-seq | T36 | 28,499 | |
| 3-Hippocampus | Mouse | Slide-seqV2 | Part of Puck_200225_08 | 21,919 | Fig. S3-4 |
| | Macaque | Stereo-seq | T36 | 28,499 | |
| 4-Hippocampus | Mouse | Stereo-seq | T315, T319, T323 | 16,256 | Fig. S5 Fig. S8 |
| | Macaque | Stereo-seq | T36, T38, T42 | 83,262 | |
| 5-Cortex | Mouse (visual cortex, VIS) | MERFISH | NA | 5,995 spots, 234 genes | Fig. S6 |
| | Human (auditory cortex, AUD) | MERFISH | NA | 3,970 spots, 4,000 genes | |
| 6-Embryo (3 species) | Human | 10x Visium | Slice 2 | 3,145 | Fig. S10, S11 Fig. S26c-26d |
| | Mouse | Stereo-seq | E11.5 | 30,124 | |
| | Zebrafish | Stereo-seq | 24hpf | 5,271 | |
| 7-Embryo (2 species, 3 stages) | Mouse | Stereo-seq | E9.5, E11.5, E12.5 | 87,402 | Fig. 4 Fig. S9 |
| | Zebrafish | Stereo-seq | 12hpf, 18hpf, 24hpf | 10,400 | |

| | | | | | |
|---|---|---|---|---|---|
| 8-PFC (cell type) | Human (PFC) | Slide-tags | NA | 12,574 | Fig. S24a-S24e Fig. S25 |
| | Macaque (DLPFC) | Stereo-seq | Part of T127 | 5,859 | |
| 9-Cerebellar (cell type) | Mouse | Stereo-seq | T170 | 35,753 | Fig. 24f, 24g Fig. S25c, 25d, 14e |
| | Marmoset | Stereo-seq | T484 | 27,390 | |
| | Macaque | Stereo-seq | T44 | 53,264 | |
| 10-Hippocampus (AD) | Mouse | Slide-seqV2 | Part of j20_rep1 | 15,092 | Fig. 6a-6e Fig. S12, S13, S14 |
| | Macaque | Stereo-seq | T36 | 28,499 | |
| 11-Breast (breast cancer) | Mouse | 10x Visium | NA | 1,978 | Fig. 6f-6k Fig. S16 |
| | Human | 10x Xenium in situ | sample1, rep1 | 160,303 | |
| | Human | 10x Xenium in situ | sample1, rep2 | 118,752 | |
| 12-Liver (liver damage) | Mouse | Stereo-seq | D17_FS3 | 27,915 | Fig. S17, S18 |
| | Human | 10x Visium | | 2,265 | |
| 13-Cerebellar (three dimensional) | Mouse | Stereo-seq | 32 coronal sections: T167, T168, T169, T170, T171, T172, T173, T174, T175, T176, T177, T178, T179, T180, T181, T182, T183, T184, T185, T186, T187, T188, T189, T190, T191, T192, T193, T194, T195, T196, T197, T198 | 1,810,619 | Fig. 7 Fig. S19-S23 |

| | | | | |
|---|---|---|---|---|
| | Marmoset | Stereo-seq | 32 coronal sections: T479, T480, T482, T483, T484, T485, T486, T487, T488, T489, T490, T491, T493, T495, T496, T497, T499, T501, T502, T503, T504, T505, T506, T509, T511, T513, T514, T515, T516, T517, T518, T519 | 1,916,578 |
| | Macaque | Stereo-seq | 34 coronal sections: T40, T42, T44, T46, T48, T50, T52, T54, T56, T58, T60, T62, T64, T66, T68, T72, T74, T76, T80, T82, T84, T86, T88, T90, T92, T94, T96, T98, T100, T102, T104, T106, T108, T110 | 8,016,404 |

**Benchmarking methods settings**

***scRNA-seq integration methods.*** The high performance of Harmony and Scanorama has been shown in a previous benchmarking study for scRNA-seq data integration. According to the standard workflow, PCA was first performed on the combined expression matrix. Then, the obtained PCA embeddings and batch information were considered as input of the harmony_integrate() function in the SCANPY package. Finally, the corrected PCA embeddings were obtained as output. The number of principal components is set to 32 to 64 for different datasets (default setting 32).

***Spatial transcriptomics integration methods.*** SEDR is a spatial embedding representation method based on graph neural networks with PCA embeddings as node features. In the original study, the authors fed the latent SEDR embeddings to Harmony for batch correction. We empirically disabled the deep embedded clustering (DEC) loss function to obtain better performance. The number of principal components is set as 300, and the epoch is set as 300 (default setting). STAligner is a graph attention neural network for integrating multiple ST datasets. For constructing the spatial neighbor graphs, we kept the hype-parameters consistent with STACAME. We optimized the training hyper-parameters for STAligner.

***Spatial cell-type annotation method.*** STELLAR is a cell-type predicting method for spatially resolved single-cell datasets that utilizes a graph convolutional neural network. We optimized the training hyper-parameters of STELLAR for each experiment.
